# Predicting the long-term impact of AI assistants on collective intelligence

**DOI:** 10.64898/2026.09.29.755292

**Authors:** Jaeseung Kim, Sang Woo Park

## Abstract

Artificial intelligence (AI) assistants can improve individual performance but may weaken unaided performance and limit interest diversity, raising questions about their long-term effects on collective intelligence (CI). Here, we consider an evolutionary model in which individuals predict, consult AI, and imitate better-performing peers. Classic reward structures that promote CI without AI can fail under “Chatbot AI”, whose predictions depend on users’ interests, by reducing interest diversity or propagating bias. Directly rewarding or penalizing AI use does not reliably mitigate these effects. Instead, a balanced incentive that preserves minority expertise and rewards collective contributions can promote CI, but an additional incentive is needed to restore unaided collective accuracy. Our framework offers an explanation for emerging evidence of AI’s societal impact and clarifies its long-term implications.

---

Collective intelligence (CI) describes a phenomenon in which a group’s ability to solve problems, make decisions, or adapt to changing environments exceeds each member’s ability (*1–5*). This phenomenon, also referred to as wisdom of crowds, has thus served as a central mechanism underlying the success of diverse organizations, ranging from animal groups (*6–8*) and human societies (*4, 9*) to machine-learning systems (*10, 11*). Understanding the conditions that promote and maintain CI therefore remains a fundamental question across social sciences (*12*), evolutionary biology (*13*), and computer science (*14*). Addressing this question has become particularly timely as artificial intelligence (AI) has begun to reshape how individuals acquire information, make decisions, and contribute to collective outcomes (*15*).

The recent emergence of generative AI and large language models (LLMs) has changed the way individuals engage in various cognitive tasks, ranging from everyday tasks (*16,17*) to scientific discovery (*18*) and creative writing (*19*). In particular, while the use of generative AI models, such as ChatGPT, has been rapidly increasing (*20, 21*), the potential long-term impact of AI assistants on human society, especially CI, remains uncertain. Some argue that AI can promote CI by enhancing individual expertise (*22*) and opinion aggregation (*23, 24*). Conversely, empirical evidence indicates that the use of AI can limit the diversity of opinions (*19, 25–27*), a key requirement for the emergence of CI (*28–31*). We expect these AI-driven changes to shape long-term social evolution because successful individuals can influence which interests, beliefs, and patterns of AI use become prevalent in a population. However, our ability to predict the long-term impact of AI assistants on human CI is hampered by the apparent discrepancy in the incentives that shape AI use. On one hand, many industrial sectors are pushing towards increased AI use, where the number of “tokens” used is often perceived as a proxy for productivity (*32, 33*). On the other hand, many academic institutions appear to discourage AI use, reflecting concerns about plagiarism, misinformation, and limited independence (*34, 35*). Such discrepancy raises a question about whether directly incentivizing or penalizing AI use will benefit or harm human CI. More broadly, previous work identified incentive structures that promote CI through social learning (*13*), but it remains unclear whether these structures remain effective in the presence of AI assistants. Thus, a key challenge is to identify robust incentive structures that retain the benefits of AI while limiting risks such as the loss of diversity in individual opinions, thereby allowing for the maintenance of CI under AI use.

To address these questions, we extend previous approaches for modeling CI, in which each individual participates in a prediction task and imitates better-performing peers (*12, 13, 36*). Previous work modeled the prediction target *Y* as a linear combination of underlying causal factors, *X*_1_, *X*_2_, …, *X*_*M*_, (Fig. 1A) and assumed that each individual makes an independent prediction from a single observed factor of their interest *e*_*A*_ and their belief 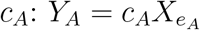 (Fig. 1B). Here, we further allow each individual to independently consult with AI assistants, which can also make their own predictions 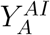, and revise their predictions by taking the weighted average according to their reliance on AI 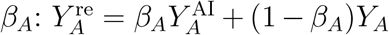 (Fig. 1B). In doing so, we consider two forms of AI: Omniscient AI constructs predictions using all causal factors, whereas Chatbot AI responds only about the factor that the user is interested in (Fig. 1B). These AI predictions can include both systematic biases and stochastic errors (Fig. 1B). Then, these revised predictions are aggregated to form a collective prediction *Ŷ*^re^, which is then used to evaluate the collective accuracy of the population (*13*): 1 − E[(*Y* − *Ŷ*^re^)^2^]*/*E[(*Y* − *α*_0_)^2^]. For Omniscient AI, we simply average revised predictions (namely, “averaging”). For Chatbot AI, we average within interests and sum across represented interests (namely, “clustering”). Hereafter, we say that CI emerges when the collective accuracy reaches a near-perfect level.

**Figure 1:**
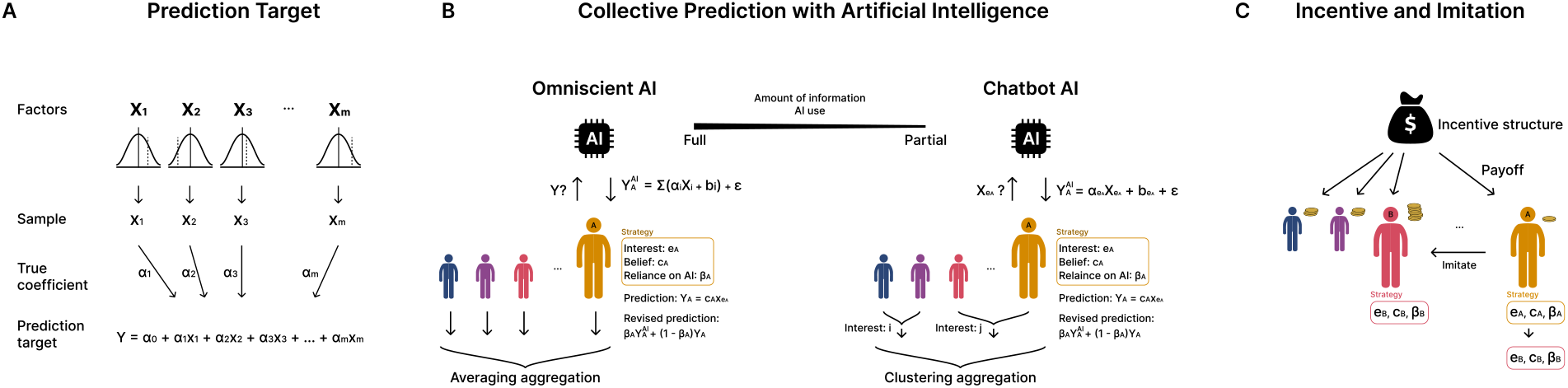
A schematic diagram of the model for individual and collective prediction in the presence of AI assistants. (A) The prediction target is modeled as a linear combination of causal factors, 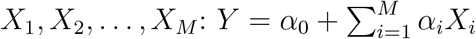. (B) Each player is characterized by three strategies: interest *e*_*A*_, belief *c*_*A*_, and reliance on AI *β*_*A*_, where where *e*_*A*_ ∈ {0, …, *M*} and *X*_0_ = 1. Players make predictions based on their interests and beliefs and subsequently revise them with AI assistance. We consider two types of AI, Chatbot AI and Omniscient AI, which differ in the amount of information available for generating predictions. Chatbot AI makes predictions based on user’s interest: 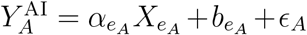. Omniscient AI makes predictions based on all causal factors: 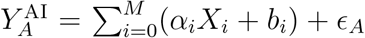, where *X*_0_ = 1. For both cases, we assume *ϵ*_*A*_ ∼ *N* (0, *τ* ). (C) Players receive payoffs according to the incentive structure, and the population evolves through imitation of peers with higher payoffs.

Individuals receive payoffs *π*_*A*_ based on their performance and imitate the interests, beliefs, and AI reliance of peers with higher expected payoffs, allowing the population to evolve over time (Fig. 1C). Following previous work (*13*), we consider two incentive structures that can promote CI in the absence of AI when paired with an appropriate aggregation rule. First, the feedback structure gives higher payoffs to individuals whose predictions are aligned with the collective prediction error, rather than directly rewarding individual predictive accuracy: 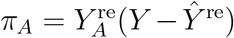. Second, the niche-expert structure gives higher payoff to individuals who have both better predictive accuracy and a minority interest: 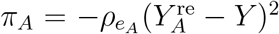, where 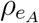 represents the proportion of individuals sharing the same interest. We assume that prediction events occur on a substantially faster timescale than social learning, which allows us to calculate the expected payoff E[*π*_*A*_] of each player analytically and simulate social learning. Full details of the model and derivations of the collective prediction, expected payoffs, and collective accuracy under the two AI models are provided in Materials and Methods as well as Supplementary Text.

We begin by characterizing the dynamics of CI with AI under two classic incentive structures from prior work (*13*). We show that the outcome depends on the information available to AI: Omniscient AI can preserve or improve collective accuracy, whereas Chatbot AI can undermine CI by limiting interest diversity or propagating systematic AI bias. Directly incentivizing or penalizing AI use fails to correct these effects and can further reduce collective accuracy. Finally, we leverage these findings to propose a balanced incentive structure that preserves minority expertise while rewarding contributions to collective accuracy, thereby restoring CI under Chatbot AI. However, even when AI-assisted CI is restored, the population’s unaided collective accuracy remains suboptimal, requiring additional incentives that explicitly reward unaided predictive ability.

## Classic incentive structures for collective intelligence can fail under Chatbot AI

First, consistent with (*13*), previously proposed incentive structures allow collective intelligence to emerge in the absence of AI (Fig. 2A,C,D). One exception is the use of simple averaging with niche-expert incentive structures, under which the collective accuracy of the population remains near zero. This is because each individual accurately predicts the contribution of their factor of interest, but averaging these partial predictions dilutes the factor-specific contributions instead of reconstructing the full prediction target (Fig. S1).

**Figure 2:**
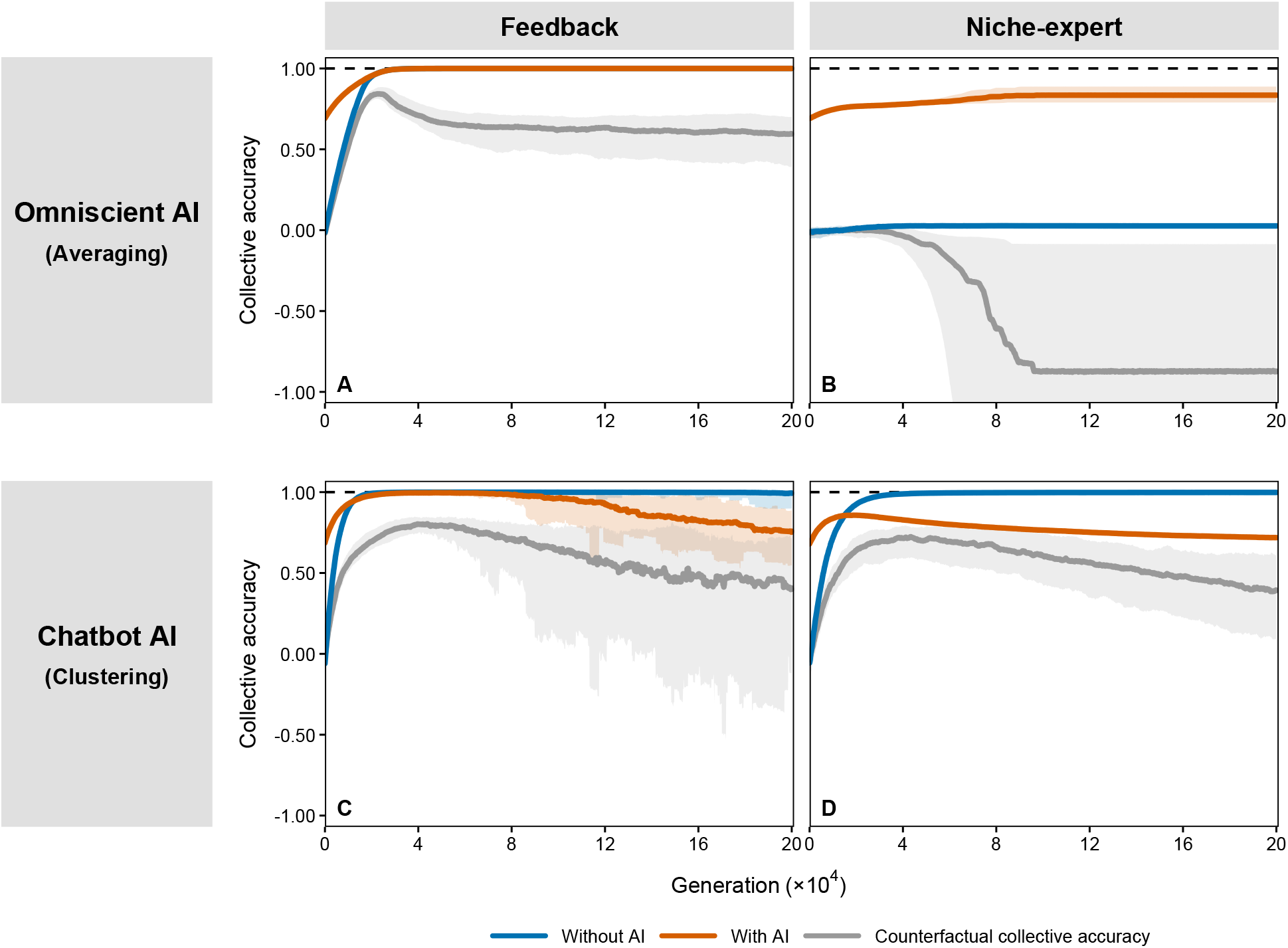
Comparison of collective accuracy with and without AI. Trajectories of collective accuracy when the group is assisted by the Omniscient AI (A,B) and Chatbot AI (C,D). Here, feedback (A,C) and niche-expert (B,D) incentive structures are considered. Blue lines represent trajectories of collective accuracy without AI assistance. Orange lines represent trajectories of collective accuracy with AI assistance. Gray lines represent trajectories of counterfactual collective accuracy, computed from players’ unrevised predictions, indicating the group’s predictive accuracy if AI assistance were unavailable at the given generation. Each condition was simulated for 30 independent replicates with randomly initialized player populations. Lines show the median across replicates, and shaded areas indicate the 95% simulation interval. Intervals extending below −1 are truncated by the y-axis limits for visual clarity. Prediction target parameters: *M* = 50, *α*_*i*_ ∼ *U* (−5, 5), and *σ*_*i*_ ∼ *U* (0, 3). Evolutionary parameters: *s* = 50 and *µ* = 0. Population parameters: *N* = 10,000, *c*_*A*_ ∼ *N* (0, 100^2^) for (A, B), *c*_*A*_ ∼ *N* (0, 5^2^) for (C, D), and *β*_*A*_ ∼ *U* (0, 1). AI parameters: *b*_*i*_ = 0.4 and *τ* = 0.3.

The use of Omniscient AI can either preserve CI under the feedback incentive structure (Fig. 2A) or promote higher collective accuracy under the niche-expert incentive structure (Fig. 2B). These results are robust across different levels of AI accuracy (Fig. S2) and can also be characterized analytically in the large-population limit (Supplementary Text).

Under the feedback incentive structure, the use of Chatbot AI leads to a loss of diversity in the population’s interest, making it impossible to account for effect of some causal factors (Fig. S2, S3). This causes an increase in collective variance and a stochastic decrease in collective accuracy (Fig. 2C; Fig. S3). We note that the loss of interest diversity may arise even in the absence of AI in finite populations (Fig. S3), but the use of Chatbot AI makes the group more vulnerable to such loss (Fig. S4). Under the niche-expert incentive structure, the population becomes overly reliant on AI, causing the collective prediction to inherit the AI’s bias (Fig. S3). This leads to an increase in collective bias and a decrease in collective accuracy (Fig. 2; Fig. S3). Interestingly, the niche-expert structure can instead reduce reliance on AI when the target intercept, *α*_0_, and AI bias have opposite signs (Figs. S2,S5). In this case, the population reduces its reliance on AI assistance, allowing CI to emerge eventually. However, the emergence of CI can be substantially delayed compared with the population without AI (Fig. S5).

To investigate whether the group can retain its predictive ability when AI assistance suddenly becomes unavailable, we quantify counterfactual collective accuracy based on unrevised predictions, retaining the population’s evolved interests and beliefs. Across all scenarios, counterfactual collective accuracy is lower than the accuracy of the corresponding AI-assisted population and the accuracy of populations that evolved without AI (Fig. 2). Thus, the accuracy achieved with AI assistance does not necessarily reflect the population’s ability to make equally accurate predictions without it.

## Direct incentives and penalties for AI do not reliably improve collective accuracy

Given the strong influence of AI on collective outcomes, a straightforward intervention is to directly incentivize or penalize the use of AI. To investigate whether such direct incentives or penalties can promote CI, we modify the two incentive structures by adding a payoff term proportional to individual reliance on AI:

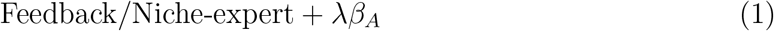

where *λ* represents the strength of incentive (*λ >* 0) or penalty (*λ <* 0), and *β*_*A*_ represents individual reliance on AI (Supplementary Text).

We find that direct incentives and penalties on AI use do not consistently improve collective accuracy (Fig. 3). When Omniscient AI is used under the feedback structure, the population retains collective intelligence regardless of the intervention strength (Fig. 3A), despite large changes in reliance on AI (Fig. 3B). In other scenarios, directly incentivizing or penalizing AI use causes individuals to imitate others based on their AI use rather than their predictive accuracy (Fig. 3C,E,G). When AI bias is relatively large, a weak penalty can lead to higher collective accuracy, but this improvement is restricted to a narrow parameter regime (Fig. 3G). Overall, these results indicate that directly rewarding or penalizing AI use may exacerbate rather than mitigate its effects on collective accuracy.

**Figure 3:**
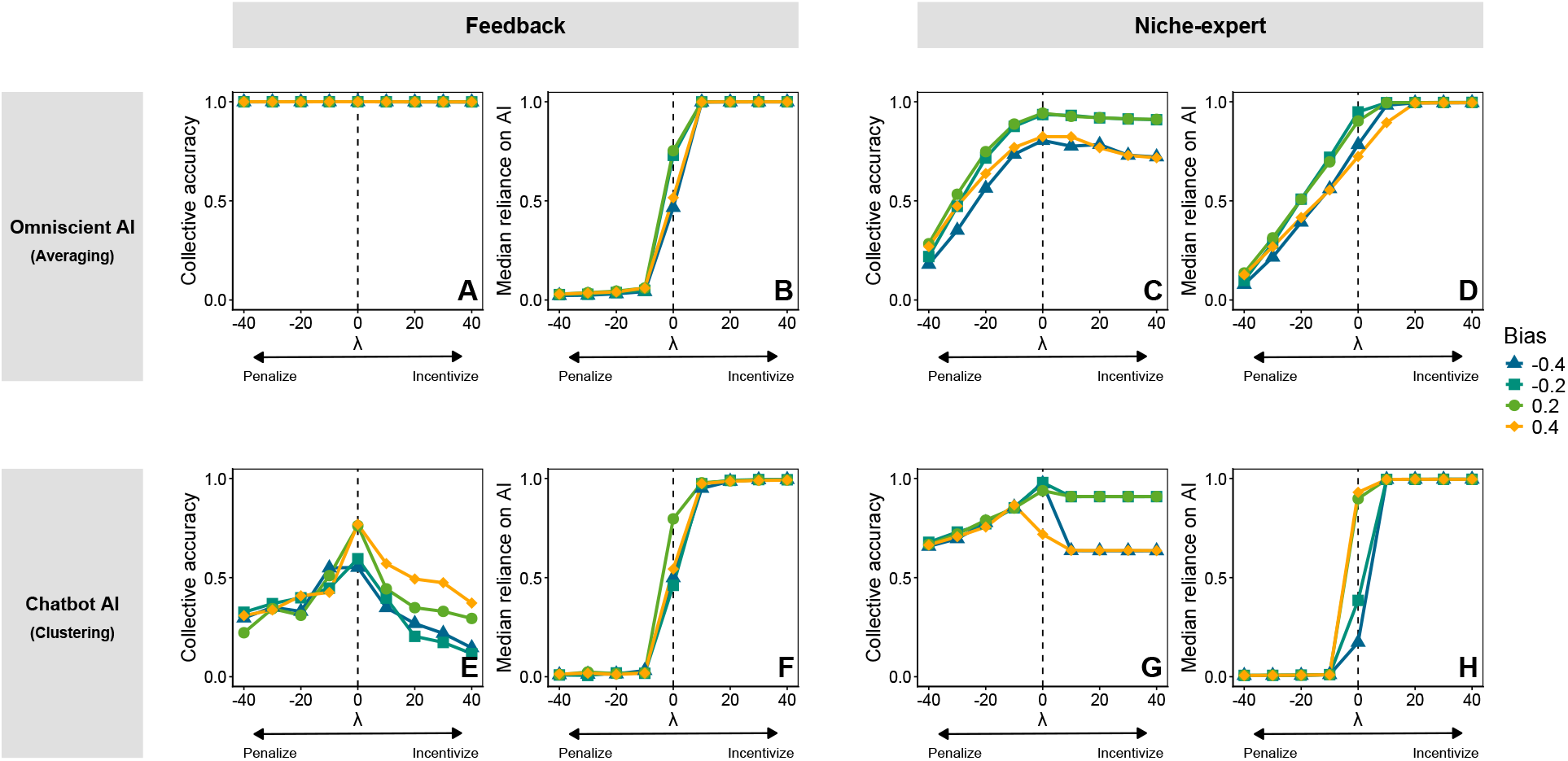
Collective accuracy under the influence of direct intervention on AI use. Collective accuracy and median reliance on AI under varying strengths of intervention (*λ*) across different AI types (Omniscient AI, A–D and Chatbot AI, E–H) and incentive structures (Feedback, A,B,E,F, and Niche expert, C,D,G,H). Positive *λ* represents an incentive for AI use, whereas negative *λ* represents a penalty for AI use. Four AI bias levels are shown: *b*_*i*_ = *±*0.2, representing higher-accuracy AI, and *b*_*i*_ = *±*0.4, representing lower-accuracy AI. Each condition was simulated for 30 independent replicates with randomly initialized player populations. Collective accuracy (A,C,E,G) and median reliance on AI (B,D,F,H) are averaged over generations 190,001–200,000 within each replicate, and points represent the median across replicates. Prediction target parameters: *M* = 50, *α*_*i*_ ∼ *U* (−5, 5), and *σ*_*i*_ ∼ *U* (0, 3). Evolutionary parameters: *s* = 50 and *µ* = 0. Population parameters: *N* = 10,000, *c*_*A*_ ∼ *N* (0, 100^2^) for (A–D), *c*_*A*_ ∼ *N* (0, 5^2^) for (E–H), and *β*_*A*_ ∼ *U* (0, 1). AI parameter: *τ* = 0.3.

## Balanced incentives promote collective intelligence with AI

The contrasting failures of the feedback and niche-expert structures suggest combining their complementary roles. Here, we propose a “balanced” incentive structure, modeled as the average of the feedback and niche expert incentives (Supplementary Text).

The core idea behind the balanced structure is to maintain interest diversity and prevent collective bias. On one hand, the feedback incentive structure is prone to the loss of interest diversity but can prevent collective bias by rewarding predictions aligned with the collective prediction error. On the other hand, the niche-expert structure can maintain interest diversity because it directly incentivizes minority interests. However, it does not directly reward the correction of collective bias, because individuals sharing the same interest are rewarded according to their individual predictive performance.

Therefore, when the two incentive structures are combined, the population achieves CI by maintaining interest diversity while limiting collective bias (Fig. 4A; Fig. S6). Under the balanced structure, individuals remain more evenly distributed across causal factors, and their beliefs lie closer to the true coefficients than under either incentive structure alone (Fig. 4C–E). The balanced structure also produces higher counterfactual collective accuracy than the feedback or niche-expert structure (Fig. 4B; Fig. S6). These patterns are robust to balancing weights (Fig. S7), and are driven by the emergent property of the balanced structure, under which the population increases its reliance on AI as AI becomes more accurate (Fig S8).

**Figure 4:**
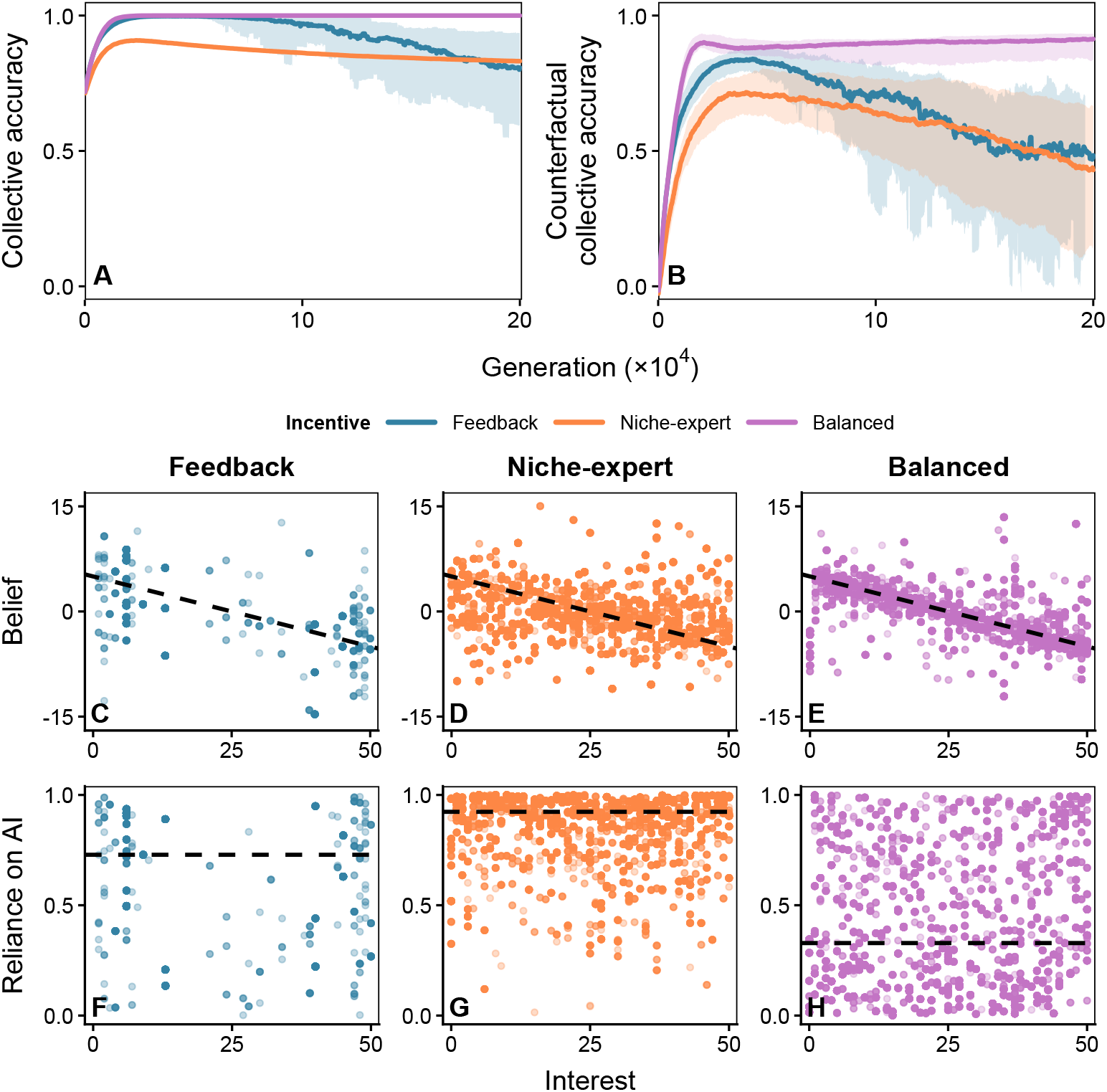
Balanced structure can promote collective intelligence. Trajectories of collective accuracy (A) and counterfactual collective accuracy (B) under three incentive structures (feedback, niche-expert, and balanced) with Chatbot AI. Each incentive structure was simulated for 30 independent replicates with randomly initialized player populations. Lines show the median across replicates, and shaded areas indicate the 95% simulation interval. Distributions of individual interest and belief (C-E) and of interest and reliance on AI (F-H) after 200,000 generations in one representative replicate. Each point represents an individual. Dashed lines in panels C-E represent the true coefficients (*α*_*i*_) of the prediction target. Dashed horizontal lines in panels F-H indicate the median reliance on AI. Prediction target parameters: *M* = 50, *α*_*i*_ = 5 − 0.2*i*, and *σ*_*i*_ ∼ *U* (0, 3). Evolutionary parameters: *s* = 50 and *µ* = 0. Population parameters: *N* = 10,000, *c*_*A*_ ∼ *N* (0, 5^2^), and *β*_*A*_ ∼ *U* (0, 1). AI parameters: *b*_*i*_ = 0.4 and *τ* = 0.3.

However, counterfactual collective accuracy remains suboptimal even under the balanced structure because payoffs are based only on revised predictions. An analogous incentive based on unrevised predictions raises counterfactual collective accuracy to near-perfect levels but does not calibrate reliance on AI, causing collective accuracy to decline as the magnitude of AI bias increases. Combining the two structures avoids this trade-off and maintains near-perfect collective accuracy both with and without AI across the range of AI bias examined (Supplementary Text; Fig. S9).

## Discussion

The ongoing race towards the development of “better” AI continues to pose questions about its societal benefits and risks (*15,23,37*). Here, we used mathematical models of evolutionary dynamics to investigate how AI use and incentive structures jointly affect the emergence of CI. We find that an “Omniscient” AI model, which incorporates information about all relevant causal factors, can improve collective accuracy, whereas a “Chatbot” AI model, whose outcome depends on the user’s interest, can impair it. These contrasting outcomes arise because AI assistance affects not only individual predictions, but also the population’s interest diversity, beliefs, and reliance on AI. We also show that directly incentivizing or penalizing AI use, independent of individual’s performances, can further limit the collective accuracy of the population. Instead, a balanced incentive structure can maintain CI by promoting diversity and limiting bias.

Our analysis reveals that AI can prevent the emergence of CI through two different mechanisms. First, AI use can cause individuals to concentrate on a subset of the factors that determine the prediction target, thereby reducing diversity in their interests. This result is broadly consistent with recent empirical studies showing that AI assistance can improve individual performance but limit the diversity of outputs, in domains ranging from scientific research (*38*) to creative writing (*19*). Second, over-reliance on AI can cause the population to inherit inherent biases in AI’s predictions (*25, 39*), causing the prediction accuracy to decrease eventually. This mechanism can prevent the emergence of CI even when the interest diversity is maintained because biases in AI predictions can be amplified when individual predictions are aggregated. Thus, preserving the diversity of ideas will not be sufficient to promote CI. Instead, AI use need to be calibrated to prevent systematic AI biases from becoming collective biases (*40*).

Our analysis of counterfactual collective accuracy also raises an important concern for future AI use: while AI use may improve the apparent performance of individuals and, in some cases, the population as a whole, it can simultaneously reduce the population’s ability to make accurate predictions without AI. These results echo findings from a recent field experiment, which demonstrated that students, who seemingly performed better in solving mathematics problems under AI assistance, performed worse on their own than those who did not have access (*41*). Our results, however, also suggest that this loss of unaided performance is not inevitable. Incorporating incentives based on individuals’ predictions without AI assistance can improve counterfactual collective accuracy without losing overall collective accuracy. These findings highlight the need to evaluate AI tools not only by the performance gains they provide during use, but also by whether individuals and populations retain the ability to perform without them (*42, 43*). Making this distinction clear will be particularly important for education, where AI may help students complete a task without helping them learn how to do it on their own (*41, 42*).

As generative AI and large language models are already generating substantial benefits, many organizations, both industrial and academic, are increasingly seeking to improve their collective performance by regulating AI use (*32–35*). Such regulations may take direct forms, including AI-use policies or token-based leaderboards, or may operate indirectly through other incentives, such as productivity rewards or the social stigma associated with AI use (*44*). Our model suggests that incentivizing or penalizing AI use is unlikely to induce appropriately calibrated reliance on AI or to promote CI. Incentivizing AI use could lead to overreliance, causing groups to propagate AI biases, whereas penalizing AI use could lead to underreliance, preventing groups from using AI even when it performs well in the relevant domain. Our results instead show that CI and appropriate reliance on AI can be promoted by rewarding diversity and contributions to collective accuracy (*12, 28, 29*), rather than by directly rewarding or penalizing AI use.

There are several limitations to our analyses. First, our model allows for social learning among humans, but AI’s inherent ability remains fixed throughout the evolutionary process. In reality, AI can also learn from human-generated data (*45*). Such bidirectional learning could increase the similarity between human and AI errors and thereby reduce group performance (*46*). Second, we model AI errors using fixed, factor-specific additive biases and independent random errors. This assumption allows random errors to be reduced through aggregation, while systematic biases can still persist in the collective prediction. In reality, AI errors may have more complex structures, including biased estimates of factor effects and correlated errors across individuals, factors, or repeated interactions. Further empirical and theoretical work is therefore needed to understand how these error structures affect the emergence of CI. Third, in our main analyses, each AI model is paired with the aggregation rule that best matches its prediction structure, making it difficult to fully separate the effects of the AI model from those of aggregation. However, analyses of alternative pairings suggest that our primary aggregation choices are consistent with the distinct prediction structures of the two AI models, although the effects of AI assistance may also depend on how individual predictions are aggregated (Supplementary Text; Fig. S10). Finally, following established models of CI, we assume that the prediction target is a linear combination of independent causal factors. This assumption does not hold for many real-world tasks, where causal factors may be correlated or interact nonlinearly. Future studies should therefore investigate how nonlinear relationships between causal factors and prediction targets affect the interaction between AI assistance and the emergence of CI.

The rapid adoption of AI assistants is changing how individuals learn, work, and make decisions (*20, 21, 24, 43, 46*). While the immediate benefits and risks of these systems have been evaluated at the individual level (*16, 41*), our study raises a broader concern that their widespread AI use may reshape collective intelligence of human society. Our results further underline the importance of designing reward structures that preserve minority expertise and prevent AI biases from persisting across the population (*12, 13, 39, 46*). A detailed empirical understanding of how AI assistance reshapes diversity, expertise, and reliance at the population level will therefore be critical for predicting the dynamics of future of human collective intelligence.

## Author contributions

Jaeseung Kim: Conceptualization, Formal Analysis, Investigation, Methodology, Software, Visualization, Writing - Original Draft Preparation, Writing - Review & Editing. Sang Woo Park: Conceptualization, Investigation, Methodology, Supervision, Writing - Original Draft Preparation, Writing - Review & Editing.

## Data, code and materials availability

All data are stored in a publicly available GitHub repository (https://github.com/parklab-snu/collective-intelligence-with-AI).

## Artificial Intelligence (AI) Disclosure

ChatGPT-5.6 was used for assistance with ggplot2 coding for figure formatting and organization.

## Supplementary materials

Materials and Methods

Supplementary Text

Figures S1 to S10

Table S1

## Materials and Methods

### Prediction target

Following previous models of CI, we model the prediction target as a linear combination of *M* independent causal factors (*12, 13, 36*):

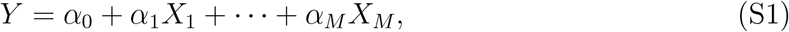

where the coefficient *α*_*i*_ determines the magnitude and direction of the effect of factor *i*, and *α*_0_ represents a global intercept. Each causal factor *X*_*i*_ is modeled as an independent Gaussian random variable with mean zero and standard deviation *σ*_*i*_.

### Players

We consider a well-mixed population *N* consisting of *N* players engaged in a prediction task (*13*). In the absence of AI assistance, the strategy of player *A* is characterized by two variables: interest *e*_*A*_ and belief *c*_*A*_. Player *A* observes the causal factor 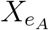 and makes a prediction by multiplying the observed factor value by their belief:

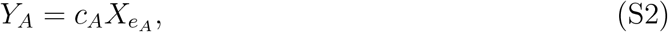

where *e*_*A*_ can take an integer value from 0 to *M* . When *e*_*A*_ = 0, the player does not observe any causal factor and instead makes a constant prediction based solely on their belief. We refer to this as a stubborn strategy because the player’s prediction does not respond to the realized values of the causal factors. For notational convenience, we assume that *X*_0_ = 1.

When the group is assisted by AI, each player’s strategy additionally includes reliance on AI, denoted by *β*_*A*_. We assume that each player consults the AI independently and revises their prediction by taking a weighted average of their own prediction and the AI-generated prediction:

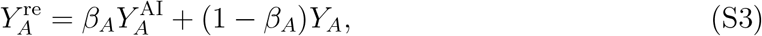

where *β*_*A*_ ranges from 0 to 1. A value of *β*_*A*_ = 1 indicates complete reliance on AI, whereas *β*_*A*_ = 0 indicates that the player does not use the AI prediction.

### AI models

We assume that the AI model has been trained on similar tasks and is therefore capable of making predictions based on causal factors. However, because the model is imperfect, its outputs are subject to systematic additive bias and stochastic error. The additive bias associated with factor *i* is denoted by *b*_*i*_, and the stochastic error is denoted by *ϵ*. The AI is neither trained nor updated during the simulation, and its biases therefore remain fixed throughout the simulation. For simplicity, we assume *b*_*i*_ = *b* for the main analysis for all *i*. The model formulation nevertheless allows biases to differ among factors.

We assume that the stochastic error follows a Gaussian distribution with mean zero and standard deviation *τ* . Each time a player consults the AI, an independent error is generated. Errors are therefore independent across players and across repeated consultations by the same player. Consequently, stochastic AI errors can be attenuated when independently generated predictions are aggregated across players. We therefore hold *τ* constant and vary the magnitude of the additive bias to manipulate the level of AI accuracy during our analyses.

We consider two extreme types of AI that differ in the amount of information available for prediction. The first is termed the Omniscient AI, which can observe all *M* causal factors and incorporate them into its prediction regardless of the user’s interest. The prediction generated by the Omniscient AI for player *A* is expressed as follows:

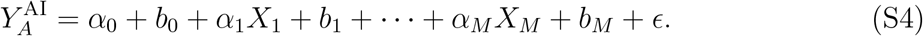

The second is termed the Chatbot AI, which can use only the single causal factor *X*_*e*_*A* associated with the player’s interest. The prediction generated by the Chatbot AI for player *A* is expressed as follows:

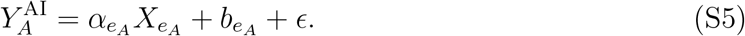

### Aggregation rules

The revised individual predictions are aggregated to form a collective prediction. We use two aggregation rules proposed in previous work (*13*), depending on the type of AI used by the group. First, when the group is assisted by the Omniscient AI, the AI component of each revised prediction contains information about all *M* factors, even though the individual’s own prediction is based on only one factor. Each revised prediction can therefore be treated as an estimate of the entire target. Accordingly, we calculate the collective prediction by taking the simple average of the revised individual predictions:

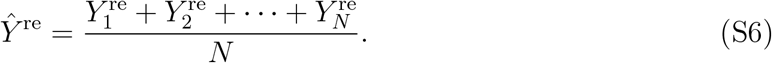

Second, when the group is assisted by the Chatbot AI, both the player’s initial prediction and the AI’s prediction are based on the same causal factor. Each revised prediction therefore contains information about only one factor and can be treated as an estimate of that factor’s contribution to the target. We consequently use clustering aggregation.

Clustering aggregation is performed by grouping individuals according to their interests and calculating the mean prediction within each group. The collective prediction is then obtained by summing the mean predictions of all nonempty groups:

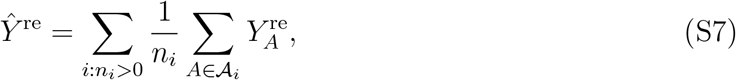

where *A*_*i*_ represents the set of players interested in factor *i*, and *n*_*i*_ = |*A*_*i*_| represents the number of players in that set.

When the group is not assisted by AI, collective predictions are computed from the unrevised individual predictions throughout the simulation. Both simple averaging and clustering can be used in the absence of AI. For fair comparison, we use the same aggregation rule as in the corresponding AI-assisted condition.

### Incentive structures

Players receive payoffs based on the underlying incentive structure. We consider seven incentive structures: feedback structure, niche-expert structure, direct intervention on AI use of these two structures, and three different balanced structures.

The feedback structure does not directly reward individuals according to their own prediction accuracy. Instead, it rewards predictions that are aligned with the residual error of the collective prediction and therefore have the potential to move the collective prediction toward the true value. Given player *A*’s revised prediction 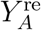 and the collective prediction *Ŷ*^re^, the payoff is expressed as

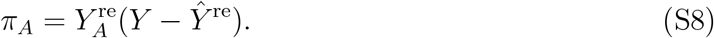

The niche-expert structure assigns higher payoffs to individuals who make accurate predictions while having an underrepresented interest. Specifically, the squared prediction error of player *A* is weighted by 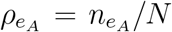, the proportion of players interested in the same factor:

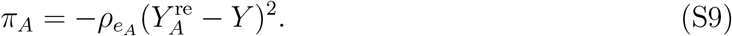

Previous research has shown that these two incentive structures can produce near perfect collective accuracy through social learning, except when the niche-expert incentive is combined with simple averaging (*13*). They are therefore used as baseline structures for examining how AI assistance affects the emergence of CI.

To examine the effects of directly encouraging or discouraging AI use, we modify the feedback and niche-expert structures by adding a term proportional to the player’s reliance on AI:

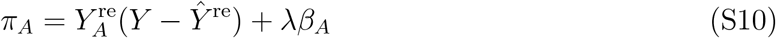

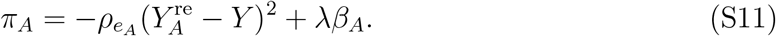

Here, *λ* represents the strength of the direct incentive or penalty applied to AI use. A positive value of *λ* rewards AI reliance, whereas a negative value penalizes it. When *λ* = 0, no direct incentive or penalty is applied, and the payoff functions reduce to the original feedback and niche-expert structures.

The balanced structure is defined as the weighted mean of the feedback and niche-expert payoff components, with balancing weight *w*:

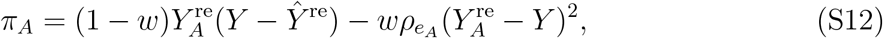

where *w* ∈ [0, 1] determines the relative contribution of the feedback and niche-expert components. We set *w* = 0.5 for all analyses unless otherwise specified, such that the two components contribute equally to the balanced structure. The sensitivity of the results to *w* is examined in Supplementary Fig. S7.

To examine whether incentives based on unrevised predictions (i.e., predictions made without incorporating AI predictions) could overcome the limitations of the other incentive structures, we defined an unrevised balanced structure. The unrevised balanced structure is defined as the arithmetic mean of the feedback structure based on the unrevised collective prediction and the niche-expert structure based on unrevised individual predictions:

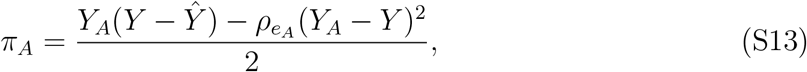

where *Ŷ* is the collective prediction based on unrevised individual predictions, and *Y*_*A*_(*Y* −*Ŷ*) and 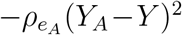 represent the unrevised feedback and unrevised niche-expert components, respectively.

We further defined a combined balanced structure as the arithmetic mean of the balanced and unrevised balanced structures, with the balancing weight set to *w* = 0.5:

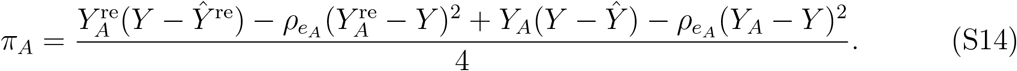

### Simulations

All simulations are conducted using an agent-based model with 50 causal factors (*M* = 50) and 10,000 players (*N* = 10,000). The parameters defining the prediction target are sampled at the beginning of the simulations and subsequently held fixed.

Coefficients *α*_*i*_ are sampled from a uniform distribution on [−5, 5], except in the simulation presented in Figure 4, in which *α*_*i*_ is set to 5 − 0.2*i* for visualization. Standard deviations *σ*_*i*_ are sampled from a uniform distribution on [0, 3].

Players’ initial strategies are also sampled from predefined probability distributions. The distribution of individual belief *c*_*A*_ depends on the aggregation rule. Under simple averaging, each factor’s contribution is scaled by the proportion of players interested in that factor. We therefore sample beliefs from a Gaussian distribution with mean zero and standard deviation 100, following the scaling used in previous work (*13*). Under clustering aggregation, beliefs are sampled from a Gaussian distribution with mean zero and standard deviation 5. Because predictions are averaged separately within each interest group and then summed, no additional scaling of *c*_*A*_ is required. The initial interest of each player is sampled uniformly from the integers between 0 and 50, and reliance on AI is sampled from a uniform distribution on [0, 1]. To account for stochastic variation, we perform 30 independent simulation replicates with randomly initialized player populations for each condition presented in the main figures.

We assume that prediction events occur on a substantially faster timescale than social learning. In each generation, we therefore calculate the expected payoff E[*π*_*A*_] of each player analytically rather than explicitly sampling prediction events. The expectation is taken over the distributions of the causal factors *X*_1_, …, *X*_*M*_ and the stochastic AI error *ϵ*, conditional on the current population strategies and the fixed target parameters. Closed form expression of expected payoffs are derived for all incentive structures and are presented in the Supplementary Text.

Each generation consists of an expected payoff calculation followed by one opportunity for pairwise social learning. Two distinct players, *A* and *B*, are randomly sampled, and player *A* imitated all three strategies of player *B* with probability

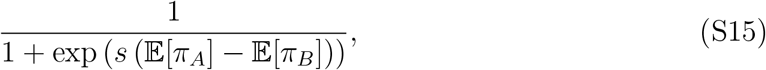

where *s* represents the strength of social learning and was set to 50 in all simulations. We introduce a mutation rate *µ*, such that with probability *µ*, player *A* does not imitate player *B* but instead redrew all strategy components from the same distributions used to initialize the population. The mutation rate is set to zero in all simulations except those presented in Supplementary Fig. S4.

Unless otherwise specified, each simulation is run for 200,000 generations. When a result is represented by a single summary value rather than as a temporal trajectory, the metric was averaged over the final 10,000 generations.

To evaluate the dynamics of collective prediction, we define six summary metrics: collective accuracy, collective bias, collective variance, counterfactual collective accuracy, interest diversity, and median reliance on AI. At each population state, expectations and variances are evaluated analytically over the distributions of the causal factors and AI errors rather than estimated from sampled prediction events. Collective bias and collective variance constitute the bias–variance decomposition of the group level collective prediction error.

Collective accuracy is normalized relative to the baseline prediction *α*_0_. A negative value indicates that the collective prediction performs worse than the baseline. Counterfactual collective accuracy is evaluated by setting *β*_*A*_ = 0 while retaining the evolved interests and beliefs of the AI-assisted population. In calculating interest diversity, we use the convention that *ρ*_*i*_ ln *ρ*_*i*_ = 0 when *ρ*_*i*_ = 0 (Table S1).

### Supplementary Text

Here, we derive the expected payoffs and collective accuracy used in the simulations. We consider Omniscient AI and Chatbot AI separately because they contain different amounts of information and therefore require different aggregation rules. We then analyze the corresponding evolutionary dynamics under a deterministic large-population approximation.

## Omniscient AI

We first consider Omniscient AI, which incorporates all causal factors regardless of the player’s interest. We derive individual and collective predictions before calculating the expected payoffs under each incentive structure.

### Model

Because Omniscient AI observes all *M* causal factors, its prediction for player *A* is given by

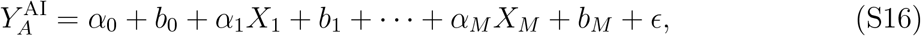

where *b*_*i*_ represent factor-specific biases and *ϵ* represents stochastic error, following a Gaussian distribution with mean zero and standard deviation *τ* . A new stochastic error is generated independently for each player and prediction event.

Let 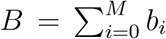 denote the total additive bias. Because the true prediction target is 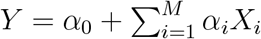, the AI prediction can be written as

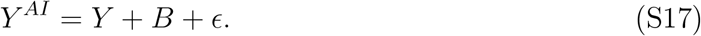

In contrast to the AI prediction, player *A*’s initial prediction depends only on the causal factor associated with their interest. Their initial and revised predictions are therefore given by:

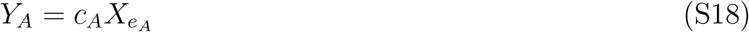

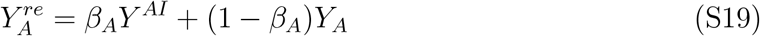

represents player *A*’s interest, *c*_*A*_ represents their belief about the effect of the corresponding factor, and *β*_*A*_ represents their reliance on AI. As in the main text, we set *X*_0_ = 1, such that a player with *e*_*A*_ = 0 makes a constant initial prediction.

### Aggregation

Because the AI component of each revised prediction contains information about all *M* causal factors, we use simple averaging to obtain the collective prediction:

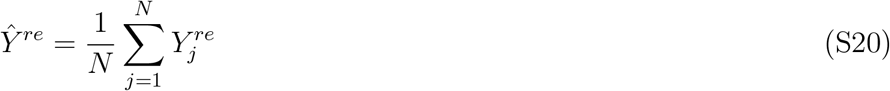

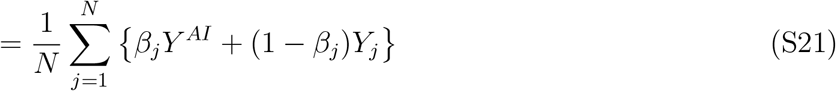

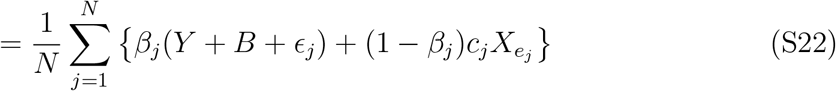

To separate the contributions of AI and the players’ initial predictions, we define the population’s mean reliance on AI, the stochastic AI error remaining after aggregation, and the aggregate contribution of initial predictions associated with each factor:

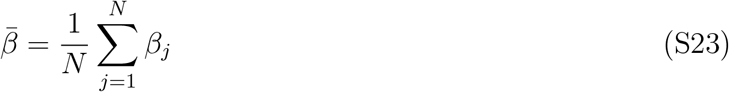

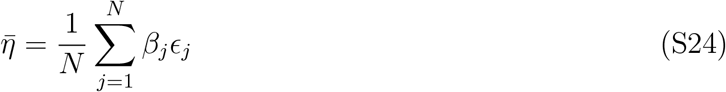

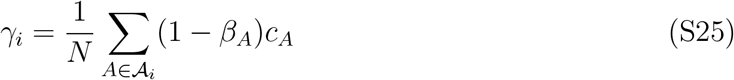

Here, 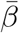 represents the mean reliance on AI, 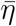 represents the stochastic AI error remaining after averaging, and *γ*_*i*_ represents the contribution of players’ initial predictions to factor *i*. Using these definitions, the revised collective prediction can be written as follows:

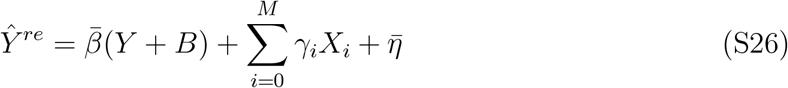

Thus, the collective prediction consists of an AI-derived component, the contributions of the players’ initial predictions, and the stochastic AI error that remains after aggregation.

### Expected payoffs

We next derive the expected payoff under each incentive structure. Expectations are taken over the distributions of the causal factors and stochastic AI errors, conditional on the current population strategies and the fixed parameters of the prediction target.

#### Feedback

Under the feedback structure, a player’s payoff depends on the alignment between their revised prediction and the residual error of the collective prediction. To express this residual, we first define the remaining error in the intercept and in the coefficient of each causal factor:

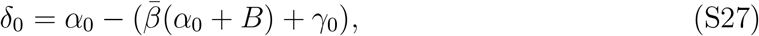

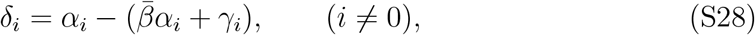

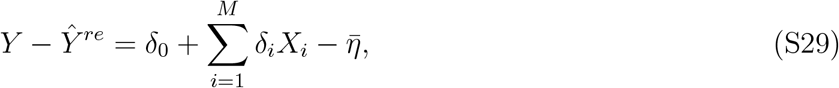

where *δ*_0_ represents the remaining error in the intercept, whereas *δ*_*i*_ represents the remaining error associated with factor *i*. Because the causal factors and stochastic AI errors are independent and have zero mean, cross terms vanish when the expectation is taken. Substituting the revised individual prediction and collective residual into the feedback payoff gives

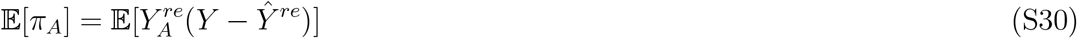

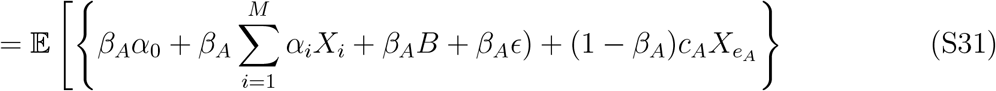

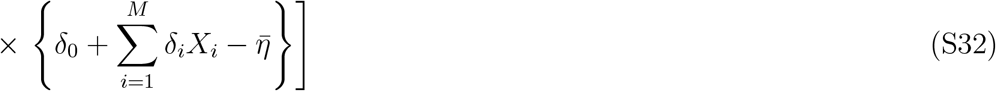

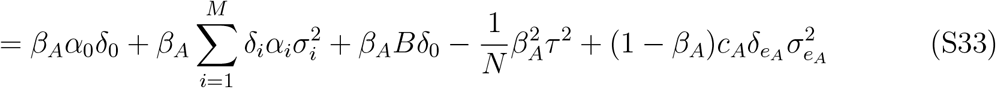

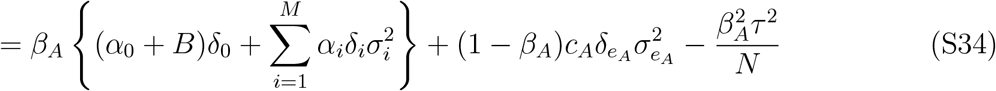

The first term represents the alignment between the AI-derived prediction and the collective residual, whereas the second term represents the contribution of the player’s initial prediction. The final term arises because the stochastic error in player *A*’s AI prediction also contributes to the collective prediction. This finite-population effect decreases as population size increases.

#### Niche-expert

Under the niche-expert structure, the payoff is the negative squared error of the revised individual prediction, weighted by the frequency of the player’s interest. Because *X*_0_ = 1, a player with *e*_*A*_ = 0 makes a constant initial prediction. In this case, the expected payoff is:

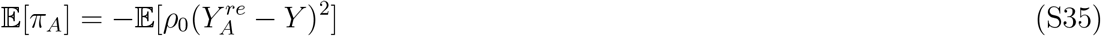

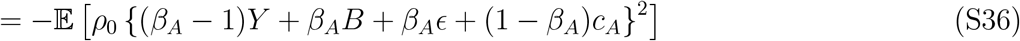

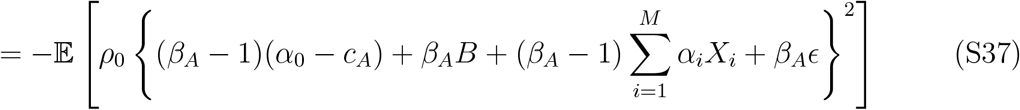

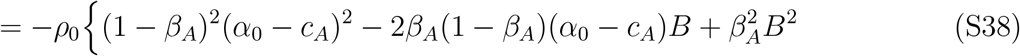

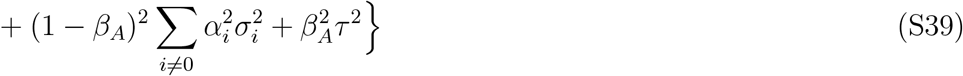

For a player with *e*_*A*_ ≠ 0, the initial prediction instead depends on the observed value of factor 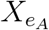 . The expected payoff is therefore:

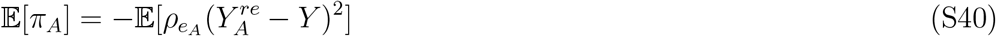

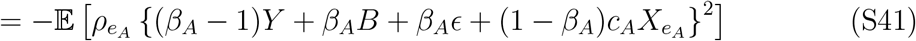

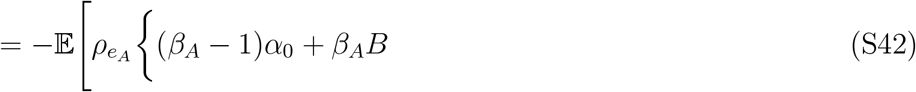

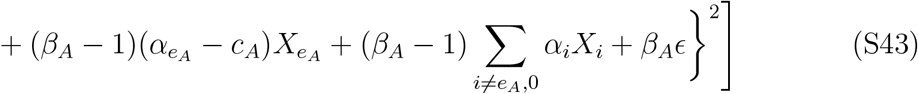

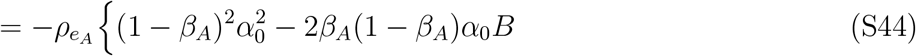

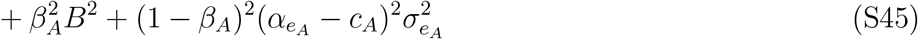

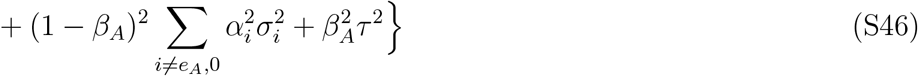

These expressions separate the contributions of the player’s belief, the unobserved causal factors, systematic AI bias, and stochastic AI error. Increasing reliance on AI reduces the contribution of errors in the player’s initial prediction but increases their exposure to AI bias and stochastic error.

#### Incentivizing or penalizing AI use

Direct incentives add the term *λβ*_*A*_ to either the feedback or niche-expert payoff. Because this term is fixed conditional on the player’s strategy, its expectation is simply *λβ*_*A*_. The expected payoffs therefore become

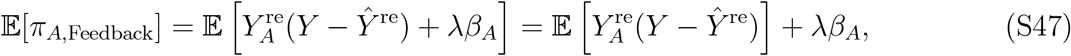

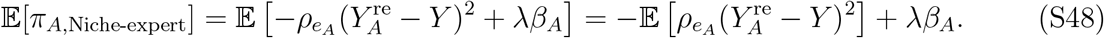

Thus, a positive value of *λ* increases the payoff of strategies with greater reliance on AI, whereas a negative value favors strategies with lower reliance.

### Expected accuracy

Using the residual defined above, the mean squared error of the revised collective prediction is:

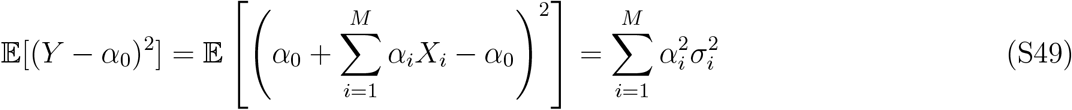

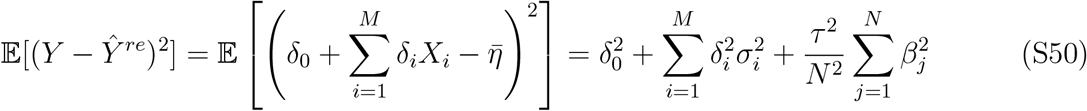

The expected collective accuracy is therefore

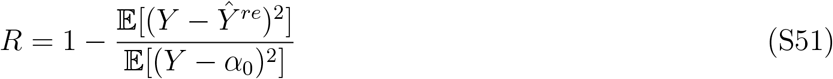

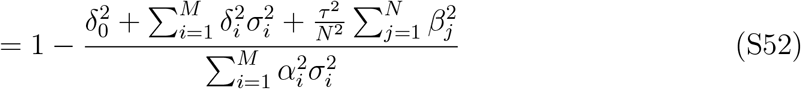

The collective prediction error consists of the squared error in the intercept, the remaining error associated with each causal factor, and the stochastic AI error remaining after averaging across players. Counterfactual collective accuracy is obtained from the same expression by setting *β*_*A*_ = 0 for all players while retaining their evolved interests and beliefs.

## Chatbot AI

Unlike Omniscient AI, Chatbot AI use only the causal factor associated with each player’s interest. This causes the information contained in AI predictions to depend directly on the distribution of interests in the population. Here, we derive the collective prediction, expected payoffs, and expected collective accuracy under this form of AI assistance.

### Model

The prediction generated by Chatbot AI for player *A* depends on the causal factor associated with the player’s interest:

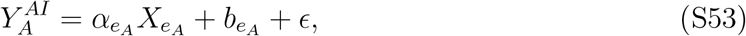

where 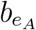 represents factor-specific additive bias, and *ϵ* represents stochastic error. The stochastic error follows a Gaussian distribution with mean zero and standard deviation *τ* and is generated independently across players and prediction events. Player A’s initial and revised predictions can be then written as follows:

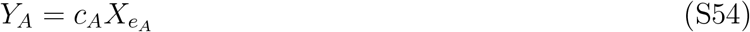

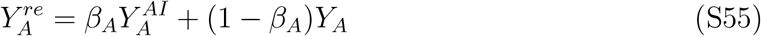

Here, *c*_*A*_ represents player *A*’s belief about the effect of their focal factor, *e*_*A*_ represents their interest, and *β*_*A*_ represents their reliance on AI. Thus, the revised prediction remains specific to the single factor associated with the player’s interest.

### Aggregation

Because both the player’s initial prediction and the Chatbot AI prediction contain information about the same causal factor, we use clustering aggregation. Revised predictions are first averaged among players interested in the same factor and then summed across represented interests:

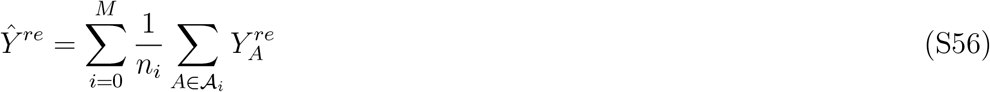

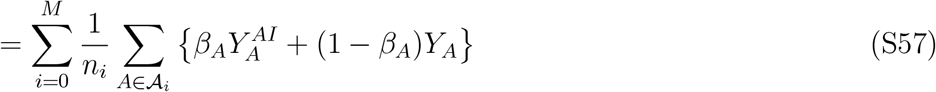

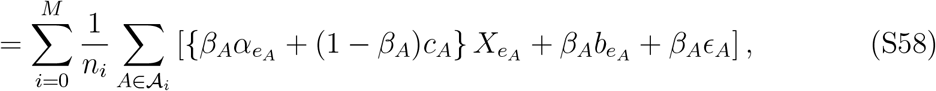

where *A*_*i*_ is the set of players interested in factor *i*, and *n*_*i*_ = |*A*_*i*_| is the number of players in that set.

To simplify the collective prediction, we define the effective coefficient, weighted AI bias, and stochastic error associated with each interest group:

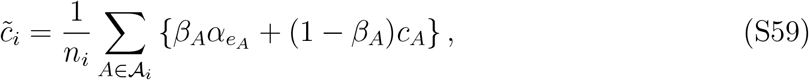

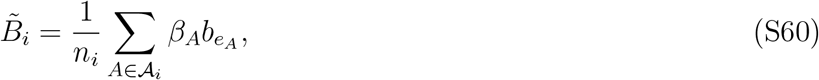

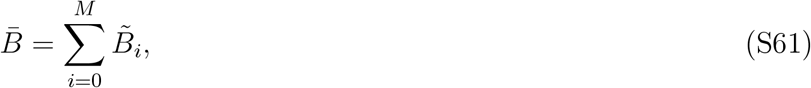

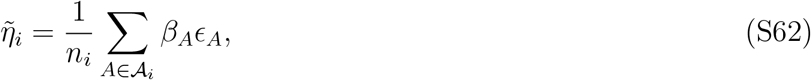

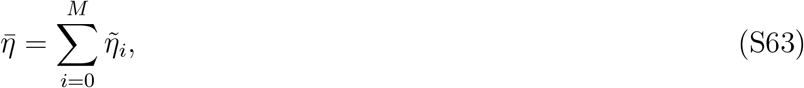

where 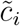 represents the mean effective coefficient for factor *i* after AI assistance, whereas 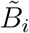and 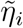 represent its systematic and stochastic error components. Thus, the collective prediction can be written as:

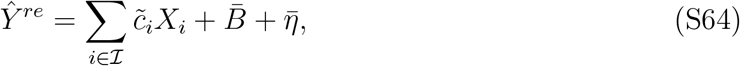

where ℐ = {*i* ∈ {0, …, *M*}|*n*_*i*_ *>* 0} denotes the set of represented interests.

### Expected payoffs

We next derive the expected payoff under each incentive structure. Expectations are taken over the causal factors and stochastic AI errors, conditional on the current population strategies and the fixed parameters of the prediction target.

#### Feedback

Under the feedback structure, players are rewarded according to whether their revised predictions reduce the error in the collective prediction. We therefore define the remaining errors in the intercept and factor-specific coefficients as

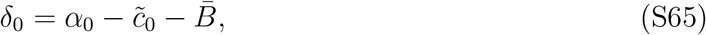

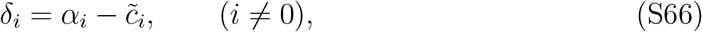

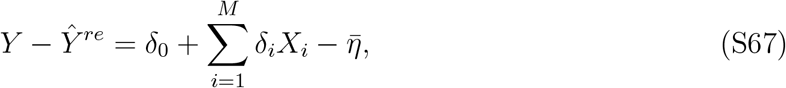

where *δ*_0_ includes the remaining error in the intercept and the total additive AI bias, whereas *δ*_*i*_ measures the remaining error in the coefficient of factor *i*. Because the causal factors and stochastic errors are independent and have zero mean, substituting the revised individual prediction into the feedback payoff gives

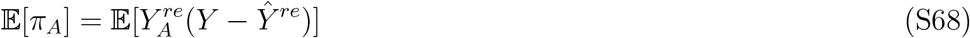

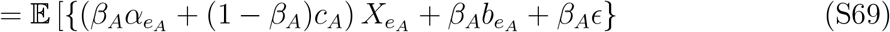

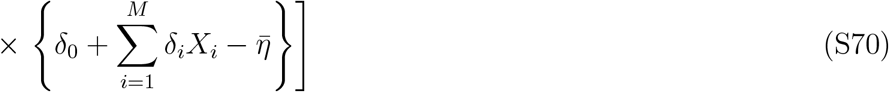

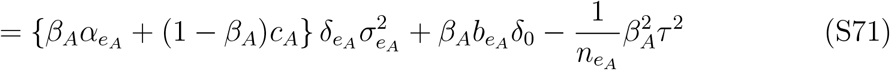

Thus, the expected feedback payoff depends on the remaining collective error associated with the player’s interest, the contribution of factor-specific AI bias to the intercept error, and the player’s contribution to the stochastic error of their interest group.

#### Niche-expert

Under the niche-expert structure, the negative squared error of each revised individual prediction is weighted by the frequency of the player’s interest. We consider *e*_*A*_ = 0 separately because these players make constant predictions rather than factor-dependent predictions.

When *e*_*A*_ = 0, the expected payoff is

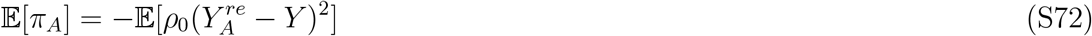

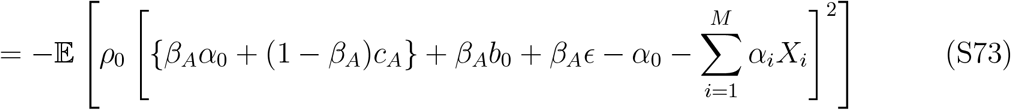

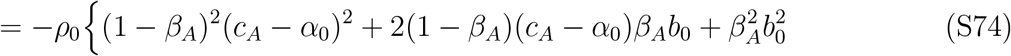

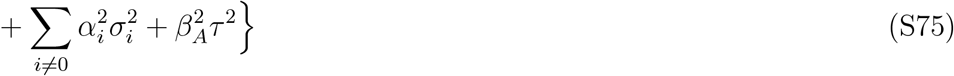

When *e*_*A*_ ≠ 0, the player’s prediction varies with their focal causal factor. The expected payoff is then

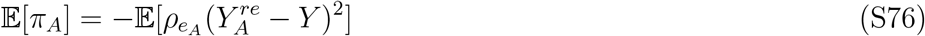

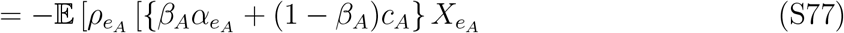

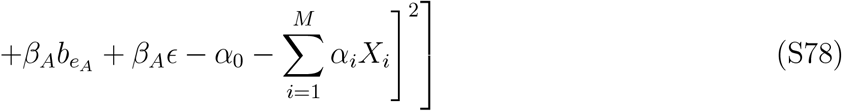

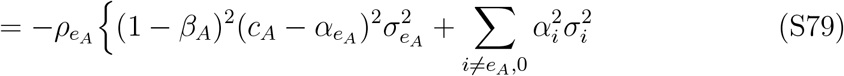

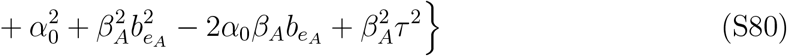

In both cases, the expected squared error consists of the mismatch between the player’s belief and the true coefficient, contributions from causal factors omitted from the individual prediction, factor-specific AI bias, and stochastic AI error. The niche-expert weight 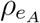 scales these errors according to the frequency of the player’s interest.

#### Incentivizing or penalizing AI use

Directly incentivizing or penalizing AI use adds the term *λβ*_*A*_ to the player’s payoff. Because this term is fixed conditional on the player’s strategy, it can be taken outside the expectation.

The expected payoffs are therefore

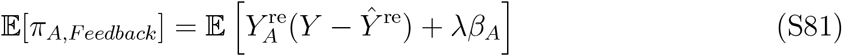

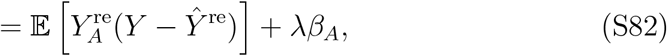

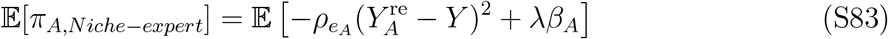

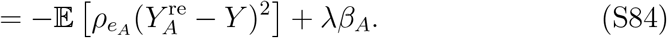

#### Balanced

Under the balanced structure, the payoff corresponds to the weighted mean of the feedback and niche-expert payoff components, with balancing weight *w*:

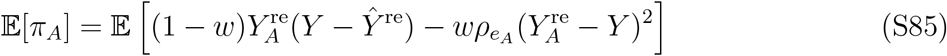

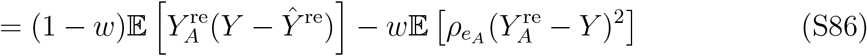

The expected payoff under the balanced structure can therefore be calculated directly from the feedback and niche-expert payoffs derived above.

#### Unrevised balanced

The unrevised feedback and niche-expert components are identical to those in the model without AI assistance (*13*). We therefore calculate their expected payoffs following the derivations in the original model and take their arithmetic mean:

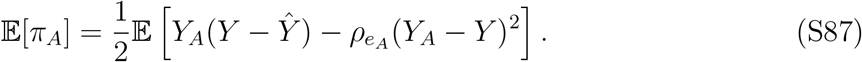

#### Combined balanced

Under the combined balanced structure, the payoff is the arithmetic mean of the balanced and unrevised balanced payoffs, with *w* = 0.5 for both balanced components. Its expected payoff is therefore

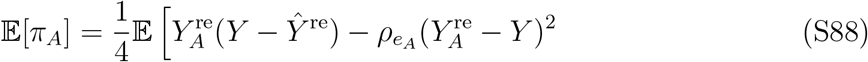

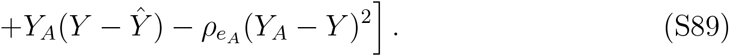

The expected payoff under the combined balanced structure can therefore be calculated directly from the revised and unrevised feedback and niche-expert payoffs derived above.

### Expected accuracy

Finally, we derive the expected collective accuracy under clustering aggregation. The intercept *α*_0_ serves as the baseline prediction. Because the causal factors are independent and have mean zero, its expected squared error is

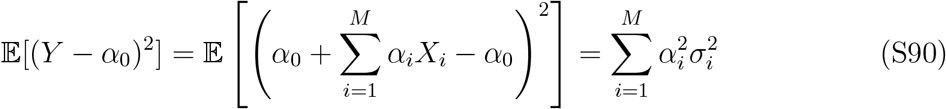

Using the expression for the collective residual above, the expected squared error of the revised collective prediction is

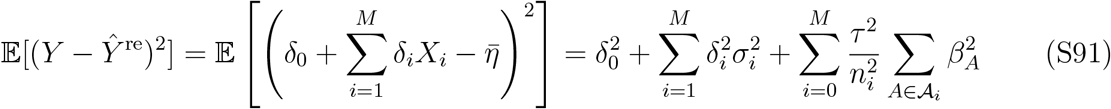

The first two terms represent squared errors in the intercept and factor-specific coefficients. The final term represents the stochastic AI error remaining after predictions are averaged within each interest group. Thus, stochastic error has a larger effect when an interest is represented by fewer individuals. Normalizing this error relative to the baseline gives:

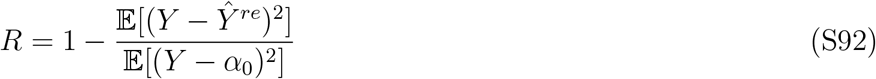

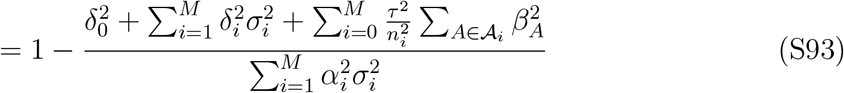

Here, *R* = 1 represents perfect collective prediction, *R* = 0 represents performance equal to the baseline, and *R <* 0 represents performance worse than the baseline.

## Evolutionary dynamics

Following previous work (*13, 47, 48*), we use replicator dynamics as a deterministic large-population approximation to characterize equilibrium states and their local stability. The agent-based simulations themselves follow the Fermi imitation rule defined above. At equal-payoff stationary states, the replicator and Fermi formulations imply the same local invasion directions for any finite selection strength *s >* 0, although their transient dynamics need not coincide. We focus on the feedback and niche-expert structures, for which the long-term dynamics can be characterized analytically. The direct interventions on AI use and the balanced structure are examined through simulations.

Let *λ* denote a strategy, *p*_*λ*_ its frequency, and *u*_*λ*_ = E(*π*_*λ*_) its expected payoff. Then, the corresponding replicator equation can be written as:

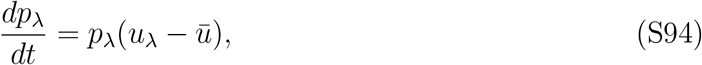

where 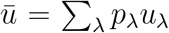 represents the mean expected payoff of the population. At an internal stationary state, all strategies with positive frequency must have the same expected payoff. A boundary state can also be stationary because a strategy with zero frequency cannot increase unless it is introduced into the population. We therefore assess the stability of boundary states by asking whether they can be invaded by a rare alternative strategy.

### Omniscient AI

With Omniscient AI, every AI prediction contains information about the full prediction target, and revised predictions are combined through averaging. As population size increases, independent AI errors vanish from the collective prediction. However, these errors remain in individual prediction errors and can therefore continue to affect niche-expert payoffs. We consider the two incentive structures separately.

#### Feedback

Under the feedback structure, both the stochastic error in the collective prediction, 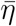, and its contribution to expected payoff vanish as *N* → ∞. Following previous work (*13*), we define the Lyapunov function as the expected error of collective prediction:

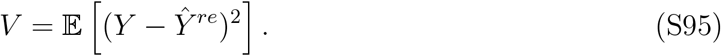

The function *V* is non-negative and equals zero when the collective prediction is identical to the target. To determine how *V* changes over time, we take its time derivative:

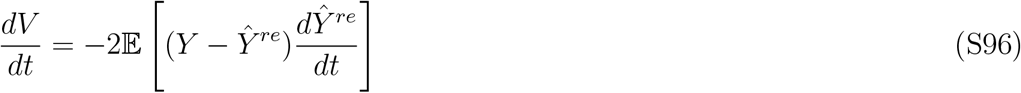

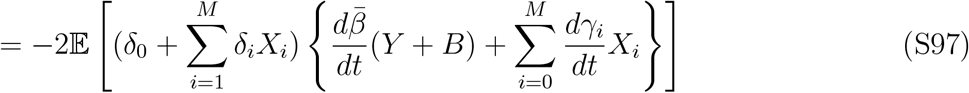

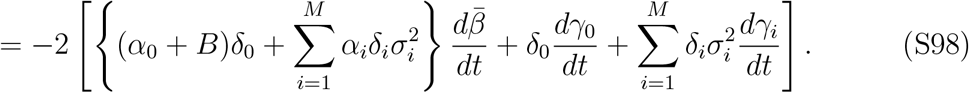

For notational convenience, we define

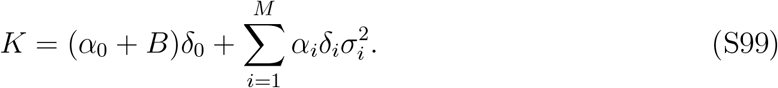

Although *K* changes with the population state, it is common to all strategies at a given state and can therefore be treated as a constant with respect to the strategy index *λ*. We also define:

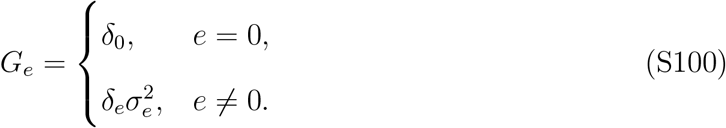

Under the replicator equation, the rate of change in the population mean of a strategy component is given by its covariance with expected payoff.

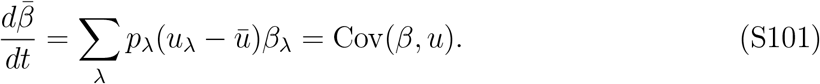

Similarly, the terms involving *γ*_*i*_ can be combined as

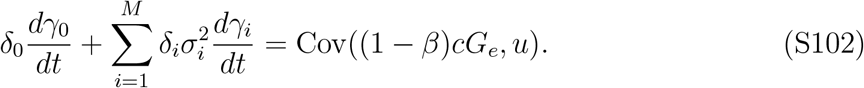

Therefore,

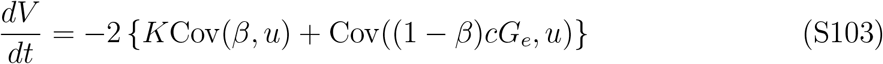

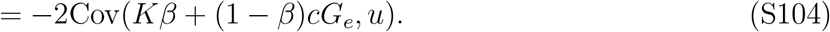

For the feedback structure in the limit *N* → ∞, the expected payoff of strategy *λ* is

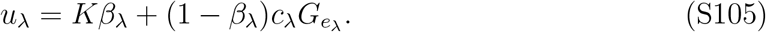

This expected payoff is identical to the quantity inside the covariance above. Therefore,

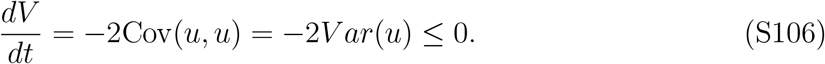

Therefore, the expected error of collective prediction cannot increase during the evolutionary process and decreases whenever strategies differ in their expected payoffs. The derivative becomes zero when the variance of expected payoffs is zero, Var(*u*) = 0. This condition arises when all represented strategies receive the same expected payoff. However, this condition does not necessarily imply that *V* = 0, because a population can lack strategies that would improve its collective prediction.

If *V >* 0, at least one component of the collective residual remains nonzero. A rare strategy whose prediction is aligned with this residual can obtain a higher expected payoff than the resident strategies and invade the population. Thus, provided that novel strategies can be introduced, an evolutionarily stable state must satisfy *V* = 0. At such a state, *Ŷ*^re^ = *Y*, and collective accuracy reaches *R* = 1.

#### Niche-expert

Unlike feedback payoffs, niche-expert payoffs depend on the prediction error of each individual. Stochastic AI error therefore remains in the expected payoff even when *N* → ∞, and we retain *τ >* 0 throughout this analysis.

We first consider strategies having the same interest by optimizing the expected payoff with respect to *c* and *β*. For a nonzero interest *e*, the expected payoff is strictly concave with respect to *c* for *β <* 1 and *σ*_*e*_ *>* 0, and is maximized at

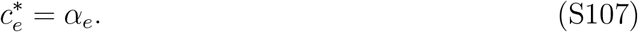

After substituting 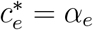, the payoff is strictly concave with respect to *β*. Its unconstrained stationary point is then:

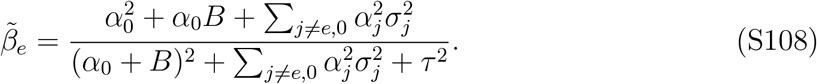

Because *β* is constrained between 0 and 1, the optimal 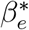 is 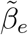 when 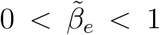, and otherwise lies at the corresponding boundary, 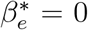 or 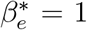. Thus, the boundary cases do not require a separate evolutionary argument; they only change the optimal prediction and its minimum expected squared error. When 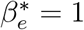, the revised prediction is independent of *c*_*e*_, so different values of *c*_*e*_ are selectively neutral and can be treated as a single payoff-equivalent strategy class.

To compare the optimized strategies across interests, let *L*_*e*_ denote the minimum expected individual squared error within interest *e*, 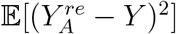. For an internal optimum,

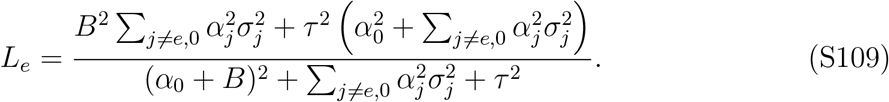

If the optimum is located at 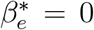 or 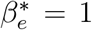, *L*_*e*_ is instead obtained by evaluating the expected squared prediction error at the corresponding boundary.

Interest 0 must be considered separately because *X*_0_ = 1 and the player’s belief determines a constant prediction. Optimizing with respect to *c* gives

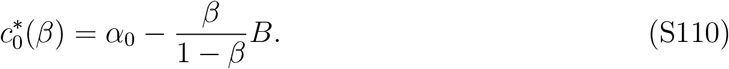

Substituting this into the expected payoff gives a strictly concave function of *β*, with the unique optimum:

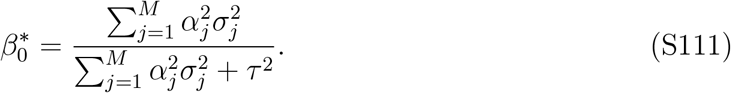

Therefore,

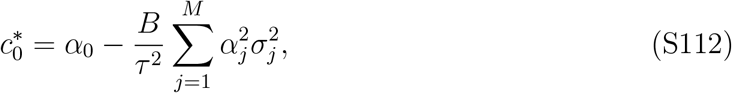

and

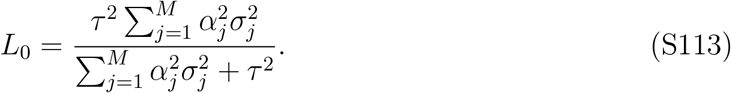

At this optimum, the player adjusts their belief to offset the systematic AI bias. The stochastic component of AI error cannot be removed in the same way and therefore limits the benefit of AI reliance.

We denote the optimal strategy within interest *e* by *S*_*e*_. When 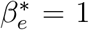, different values of *c*_*e*_ are selectively neutral and can be treated as a single payoff-equivalent strategy class. A stable state can contain only the optimal strategy class within each represented interest because a suboptimal strategy can be invaded by *S*_*e*_.

We next consider the frequencies of the *M* + 1 optimal strategies. Let *ρ*_*e*_ denote the frequency of *S*_*e*_. Its expected payoff is

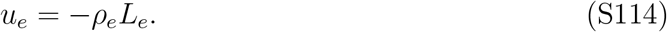

Assuming *L*_*e*_ *>* 0 for all interests, the equal-payoff condition of the replicator equation at an internal stationary state gives

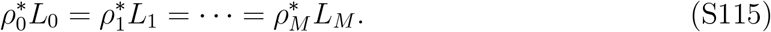

Together with 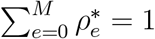, this uniquely determines

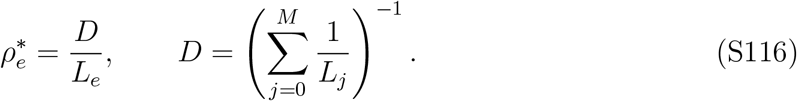

This internal stationary state is locally stable. At equilibrium, all represented interests have the same expected payoff, with 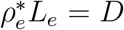. If the frequency of one interest increases slightly above its equilibrium value, its payoff decreases because the niche-expert payoff is proportional to −*ρ*_*e*_*L*_*e*_. Conversely, if the frequency of an interest decreases below its equilibrium value, its payoff increases. The replicator dynamics therefore generate a restoring force that reduces small deviations in interest frequencies and drives the population back toward the internal stationary state.

If one or more interests are absent, the resulting boundary state can be stationary under the replicator equation, but it is unstable against invasion. The payoff of a rare absent interest approaches zero as *ρ*_*e*_ → 0, whereas the mean payoff of the resident population is negative when *L*_*e*_ *>* 0, so the rare interest increases when introduced at sufficiently small frequency.

Thus, there exists a unique stable full-support equilibrium in terms of the optimal prediction strategies and their frequencies. Whether the constrained optimum has 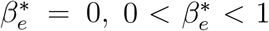, or 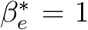 does not change the frequency argument, because these cases differ only through the resulting value of *L*_*e*_. When 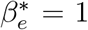, the strategy is selectively neutral with respect to *c*_*e*_, but its prediction, payoff, and equilibrium interest frequency remain uniquely determined.

Having characterized this equilibrium, we next compare its collective prediction with a prediction made entirely by AI. In particular, the expected squared error of an Omniscient AI prediction is

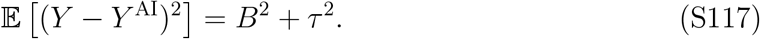

For every interest, complete reliance on AI, *β* = 1, is a feasible strategy. Therefore, because *L*_*e*_ is the minimum individual expected squared prediction error within interest *e*,

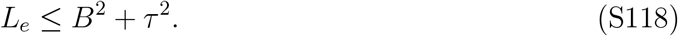

Moreover, for interest 0,

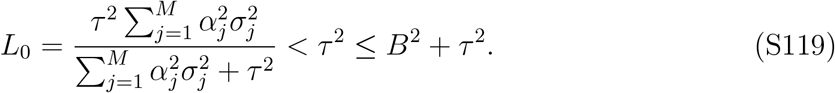

Because the stable equilibrium has 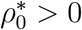,

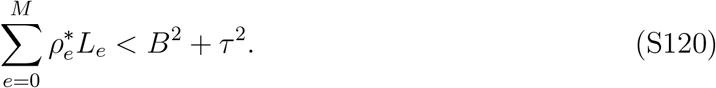

The collective prediction is the average of individual revised predictions. By convexity of the squared error,

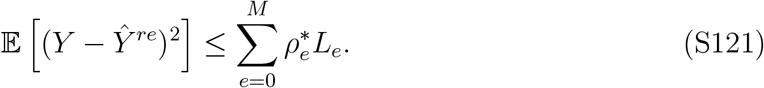

Therefore,

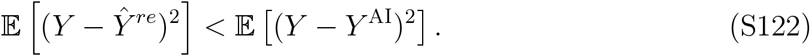

Thus, the diversity of optimized predictions allows the population to outperform an individual AI prediction at equilibrium.

However, outperforming AI does not imply perfect collective accuracy. Under averaging, the non-AI contribution associated with a particular factor is provided only by individuals interested in that factor. The predictions of individuals with other interests therefore attenuate that factor’s coefficient in the collective prediction. For a nonzero factor *i*, the residual coefficient of the stationary collective prediction can be written as

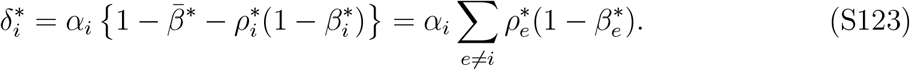

At the stable equilibrium, 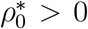. In addition, *τ >* 0 implies 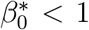. Therefore, for every factor with *α*_*i*_ ≠ 0,

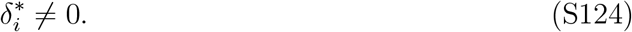

Because 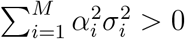, at least one factor contributes positive prediction error. Hence,

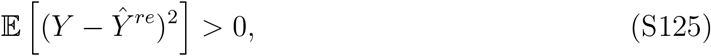

and consequently the stationary collective accuracy satisfies *R*^*∗*^ *<* 1. Thus, the niche-expert structure preserves interest diversity and allows the population to outperform AI, but averaging prevents the population from reaching perfect collective accuracy.

## AI models under alternative aggregation rules

In the main analysis, we assume that collective predictions under the omniscient AI and chatbot AI models are generated using averaging and clustering aggregation, respectively. However, the aggregation rules can, in principle, be swapped between the two AI models. To examine the consequences of these alternative combinations, we investigate their behavior through mathematical derivation and simulation.

### Omniscient AI with clustering aggregation

Under clustering aggregation, the collective prediction based on individual predictions revised by the omniscient AI can be written as

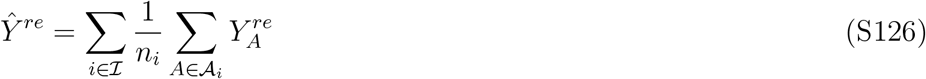

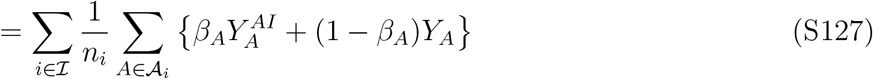

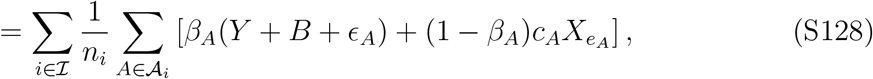

where *A*_*i*_ is the set of players interested in factor *i, n*_*i*_ = |*A*_*i*_| is the number of players in that set, and ℐ = {*i* ∈ {0, …, *M*} : *n*_*i*_ *>* 0} is the set of represented interests.

To simplify the collective prediction, we define:

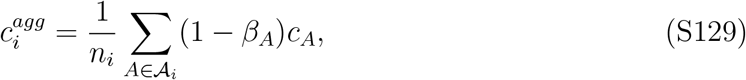

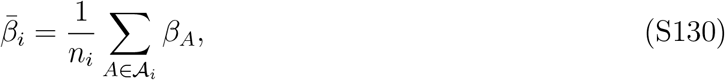

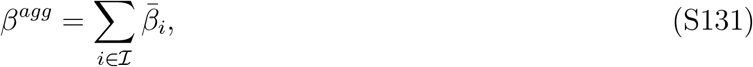

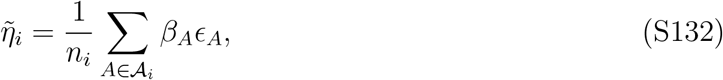

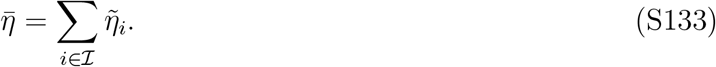

Thus, the collective prediction can be written as

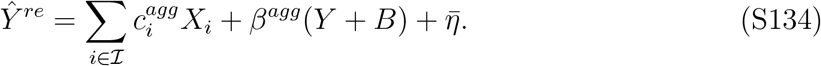

Because individual reliance on AI, *β*_*A*_, is initialized independently from *U* (0, 1), the expected initial value of *β*^*agg*^ is |*I*|*/*2. When all *M* + 1 interests are represented, this becomes (*M* + 1)*/*2. Thus, clustering aggregation incorporates the omniscient AI prediction repeatedly across interest groups, scaling its contribution to the collective prediction by a factor of approximately (*M* + 1)*/*2 at initialization. For a large number of factors, such as *M* = 50, this produces an unrealistically large contribution of the AI prediction to the collective prediction. Therefore, clustering aggregation does not provide a reasonable aggregation rule for the omniscient AI model in the context of our study, as it repeatedly incorporates predictions of the entire outcome across interest groups.

### Chatbot AI with averaging aggregation

Each prediction generated by Chatbot AI contains information about only one factor. Averaging these predictions does not repeatedly incorporate the full prediction target, but instead weights the contribution of each factor by the proportion of individuals interested in that factor. Under averaging aggregation, the collective prediction can be written as

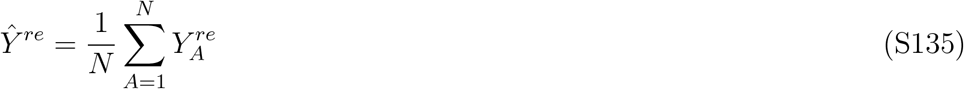

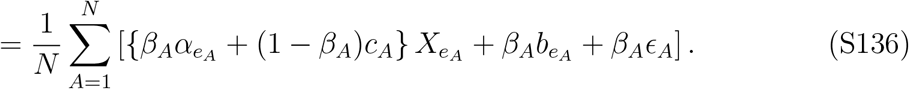

We define the aggregate revised coefficient associated with each factor, the aggregate AI bias, and the stochastic error as

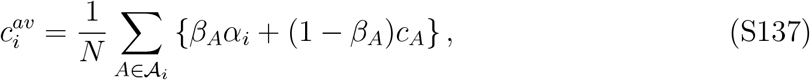

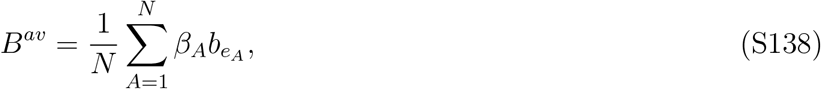

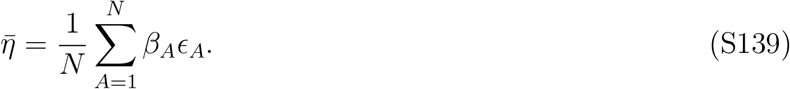

Thus, the collective prediction becomes

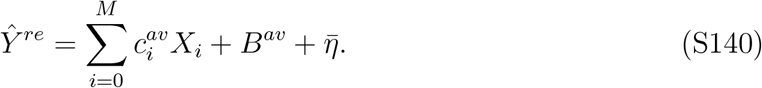

We define the remaining errors in the intercept and factor-specific coefficients as

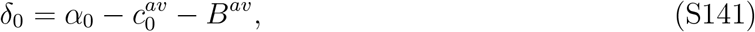

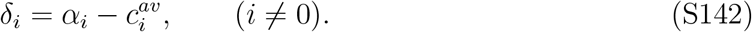

#### Expected payoffs

Under the feedback structure, the expected payoff is

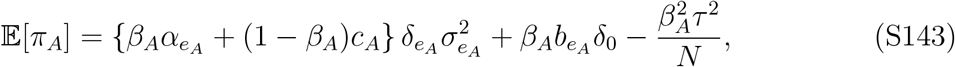

where 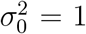. Because the niche-expert payoff depends only on individual prediction error and the frequency of the player’s interest, its expected payoff is unaffected by the aggregation rule. It is therefore identical to the expected payoff derived for Chatbot AI with clustering aggregation.

#### Expected accuracy

The expected squared error of the collective prediction is

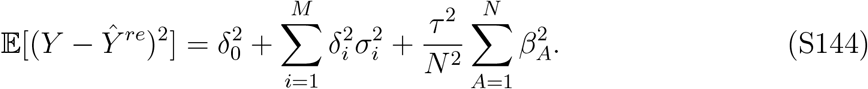

Therefore, the expected collective accuracy is

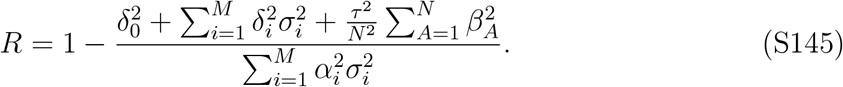

#### Simulation results

The effect of averaging depends on the incentive structure (Fig. S10). Under the feedback structure, social learning can compensate for the attenuation introduced by averaging by favoring factor-specific predictions that reduce the remaining collective error. In particular, the mean revised coefficient among individuals interested in factor *i* can approach *α*_*i*_*/ρ*_*i*_, such that its aggregate contribution approaches *α*_*i*_. Consistent with this mechanism, collective accuracy reaches a near-perfect level across AI bias values. Collective bias and variance also remain near zero, although counterfactual human CI remains below one.

In contrast, the niche-expert structure rewards individual predictive accuracy rather than contributions to the collective prediction. Revised factor-specific coefficients therefore remain close to their corresponding true coefficients, such that 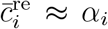. Averaging then reduces their aggregate contribution to 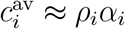. When interests are approximately uniformly distributed, *ρ*_*i*_ ≈ 1*/*(*M* + 1). The attenuation of the nonzero factor coefficients alone then gives *R* ≈ 1 − *M/*(*M* + 1)^2^ = 0.039 when *M* = 50, before accounting for the remaining intercept and stochastic errors. Accordingly, collective accuracy remains near zero even though interest diversity remains high and reliance on AI can be substantial. Chatbot AI does not resolve this problem because its predictions are also restricted to the user’s factor of interest.

These results show that the aggregation rule must reflect the information contained in individual and AI predictions. Clustering aggregation repeatedly counts the full prediction target when paired with Omniscient AI, whereas averaging aggregation attenuates factor-specific predictions when paired with Chatbot AI.

## Supplementary Figure

**Figure S1:**
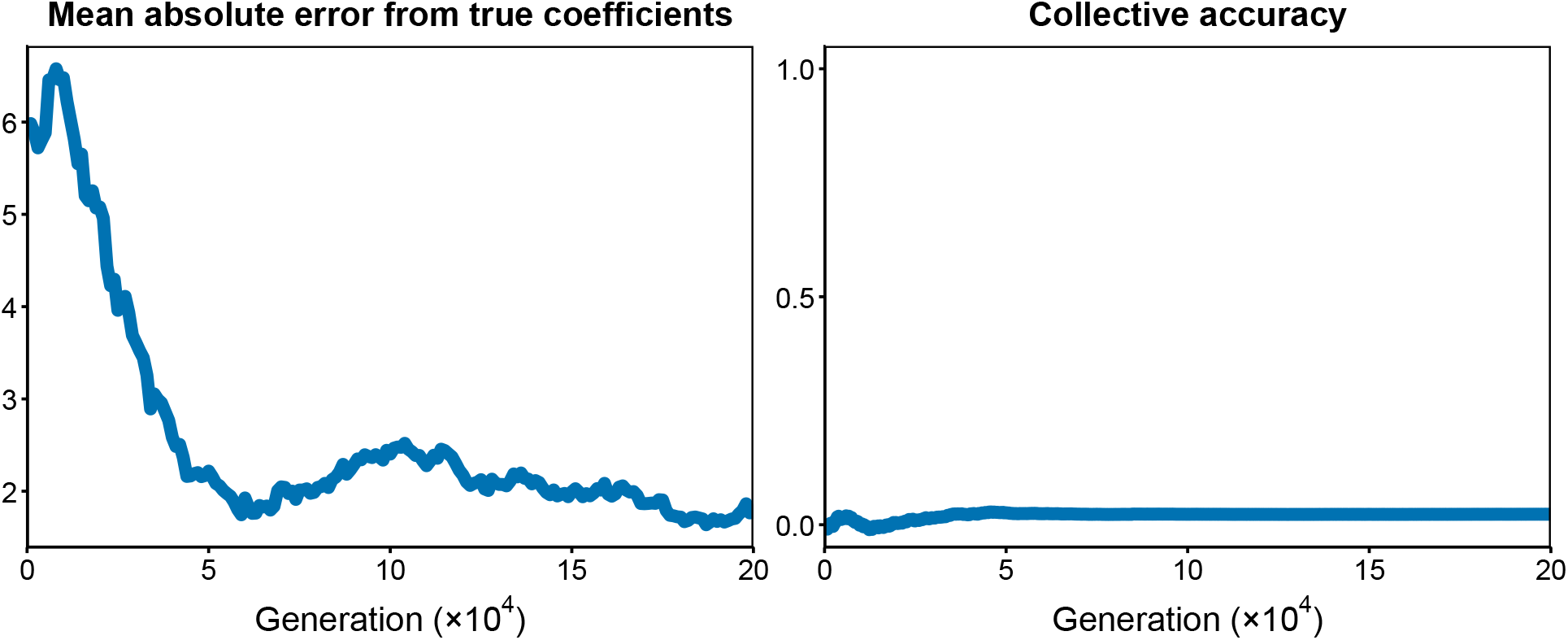
Niche-expert structure with averaging aggregation cannot promote CI without AI. Trajectories of the mean absolute error from true coefficients and collective accuracy under the niche-expert structure with averaging aggregation in the absence of AI. The mean absolute error from true coefficients is calculated by first taking, for each interest, the absolute difference between the true coefficient and the mean belief of individuals sharing that interest, and then averaging these differences across all *M* + 1 interests. The mean absolute error decreases across generations, indicating that the mean beliefs move closer to their corresponding true coefficients. However, collective accuracy remains near zero, indicating that simply averaging individual predictions does not yield accurate collective predictions. Prediction target parameters: *M* = 50, *s* = 50, *µ* = 0, *α*_*i*_ ∼ *U* (−5, 5), and *σ*_*i*_ ∼ *U* (0, 3). Population parameters: *N* = 10,000, *c*_*A*_ ∼ *N* (0, 100^2^).

**Figure S2:**
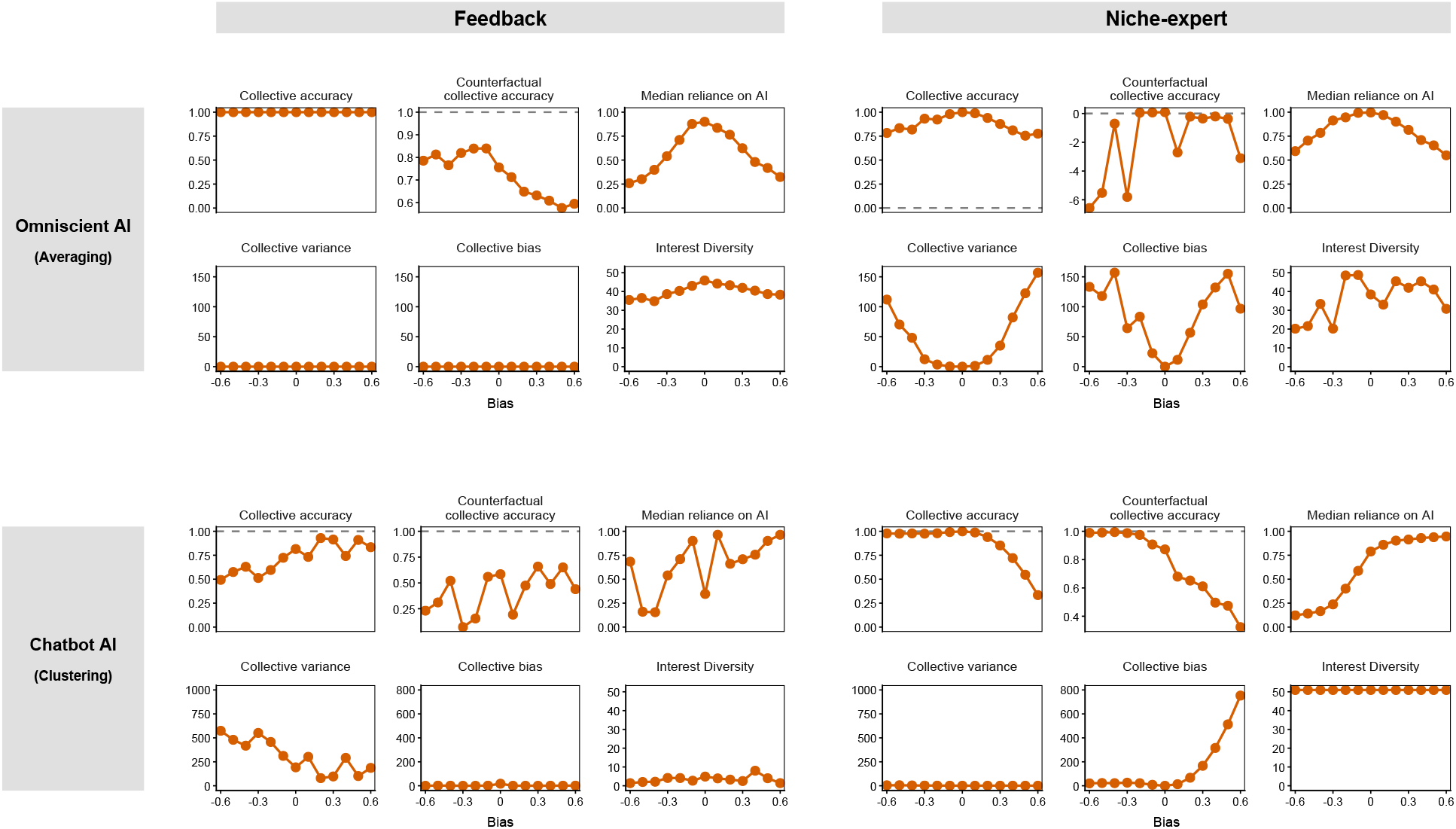
Evaluation metrics across various levels of AI bias. Evaluation metrics across varying levels of AI bias under different assumptions about AI types (Omniscient AI, and Chatbot AI) and incentive structures (feedback, niche-expert). Larger absolute AI bias indicates a less accurate AI model. Dashed horizontal lines represent the values of the evaluation metrics when the group is not assisted by AI. Collective accuracy represents the predictive performance of the group, and counterfactual human CI represents the group’s predictive accuracy if AI assistance were unavailable at the given generation. Interest diversity is defined as Hill-Shannon diversity of interest within the population, median reliance on AI represents the median value of reliance on AI (*β*). Collective bias represents systematic deviation of the collective prediction from the true target, collective variance represents variability of the collective prediction error across prediction events. All the evaluation metrics are averaged over generations 190,001–200,000. Prediction target parameters: *M* = 50, *s* = 50, *µ* = 0, *α*_*i*_ ∼ *U* (−5, 5), and *σ*_*i*_ ∼ *U* (0, 3). Population parameters: *N* = 10,000, *c*_*A*_ ∼ *N* (0, 100^2^) for (A, B), *c*_*A*_ ∼ *N* (0, 5^2^) for (C, D), and *β*_*A*_ ∼ *U* (0, 1). AI parameters: *τ* = 0.3.

**Figure S3:**
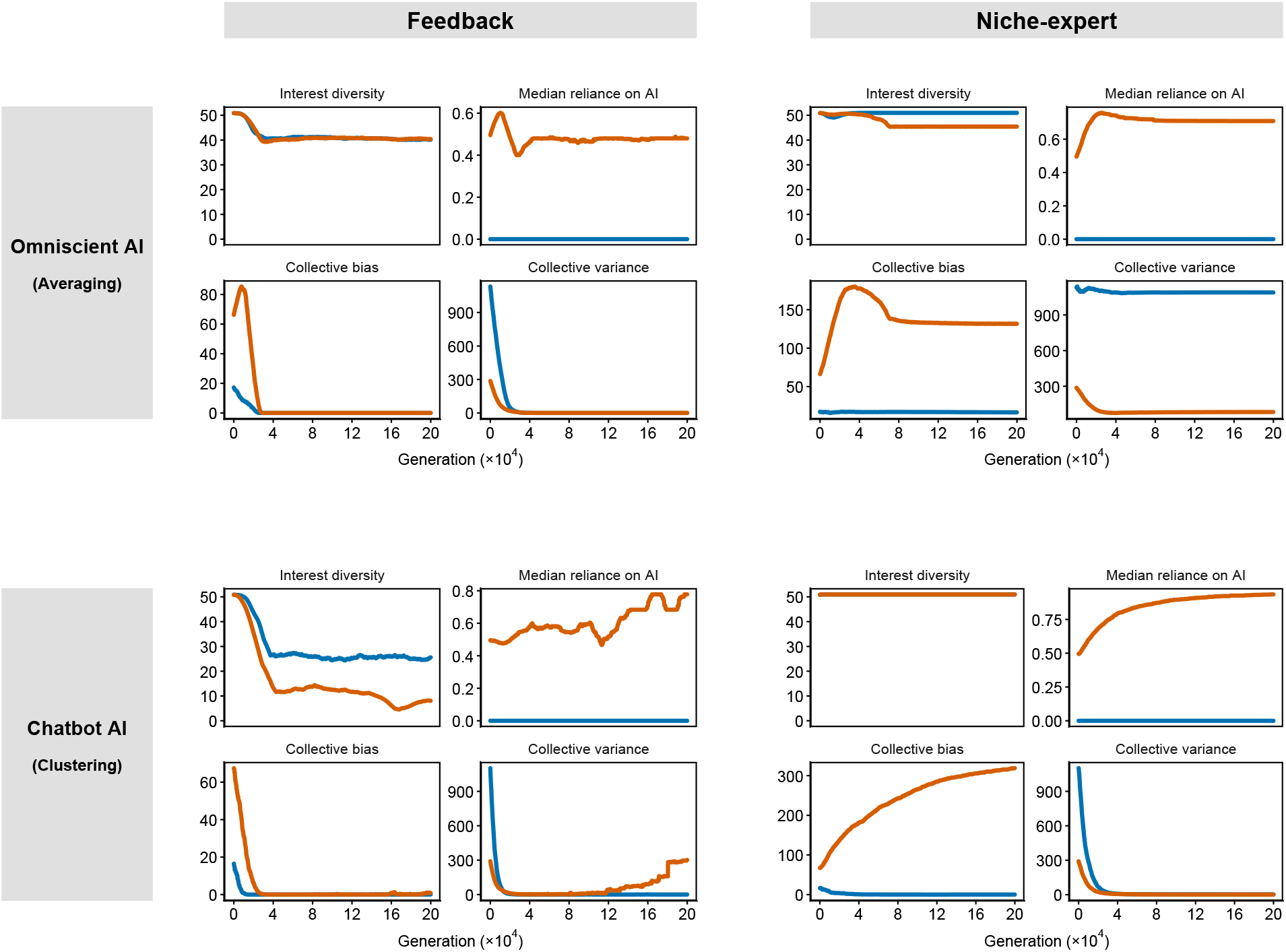
Comparison of evaluation metrics with and without AI. Trajectories of evaluation metrics when the group is assisted by the Omniscient AI and Chatbot AI. Here, feedback and niche-expert incentive structures are considered. Blue lines represent trajectories of evaluation metrics without AI assistance. Orange lines represent trajectories of evaluation metrics with AI assistance. Interest diversity is define as Hill-Shannon diversity of interest within population, median reliance on AI represent population median of reliance on AI (*β*), collective bias represent systematic deviation of the collective prediction from the true target, collective variance represent variability of the collective prediction error across prediction events. Prediction target parameters: *M* = 50, *s* = 50, *µ* = 0, *α*_*i*_ ∼ *U* (−5, 5), and *σ*_*i*_ ∼ *U* (0, 3). Population parameters: *N* = 10,000, *c*_*A*_ ∼ *N* (0, 100^2^) for (A, B), *c*_*A*_ ∼ *N* (0, 5^2^) for (C, D), and *β*_*A*_ ∼ *U* (0, 1). AI parameters: *b*_*i*_ = 0.4 and *τ* = 0.3.

**Figure S4:**
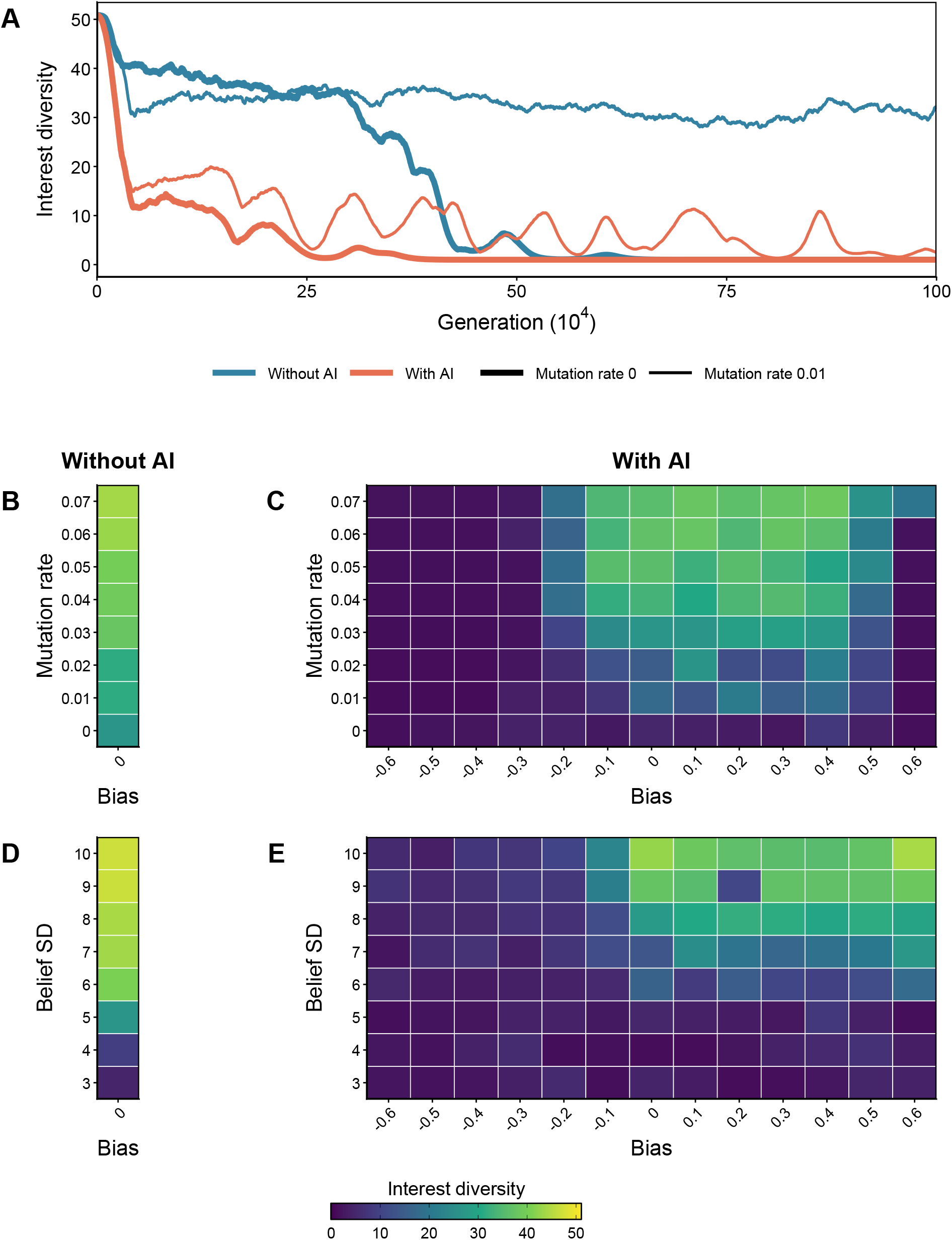
Feedback structure becomes more prone to interest diversity loss with AI. Trajectories of interest diversity with and without AI under the feedback structure (A). Heatmaps of interest diversity across varying levels of mutation rate (B, C) and the standard deviation of the initial belief distribution (D, E), with and without AI. Interest diversity declines toward zero even without AI over long timescales when there is no mutation and the initial belief distribution is narrow. However, with AI, interest diversity declines more strongly, even under moderate mutation rates and wider initial belief distributions. Interest diversity in the heatmaps is averaged over generations 190,001–200,000. Prediction target parameters: *M* = 50, *s* = 50, *µ* = 0 for (D, E), *α*_*i*_ ∼ *U* (−5, 5), and *σ*_*i*_ ∼ *U* (0, 3). Population parameters: *N* = 10,000, *c*_*A*_ ∼ *N* (0, 5^2^) for (A, B, C), and *β*_*A*_ ∼ *U* (0, 1). AI parameters: *b*_*i*_ = 0.4 and *τ* = 0.3, Chatbot AI is used.

**Figure S5:**
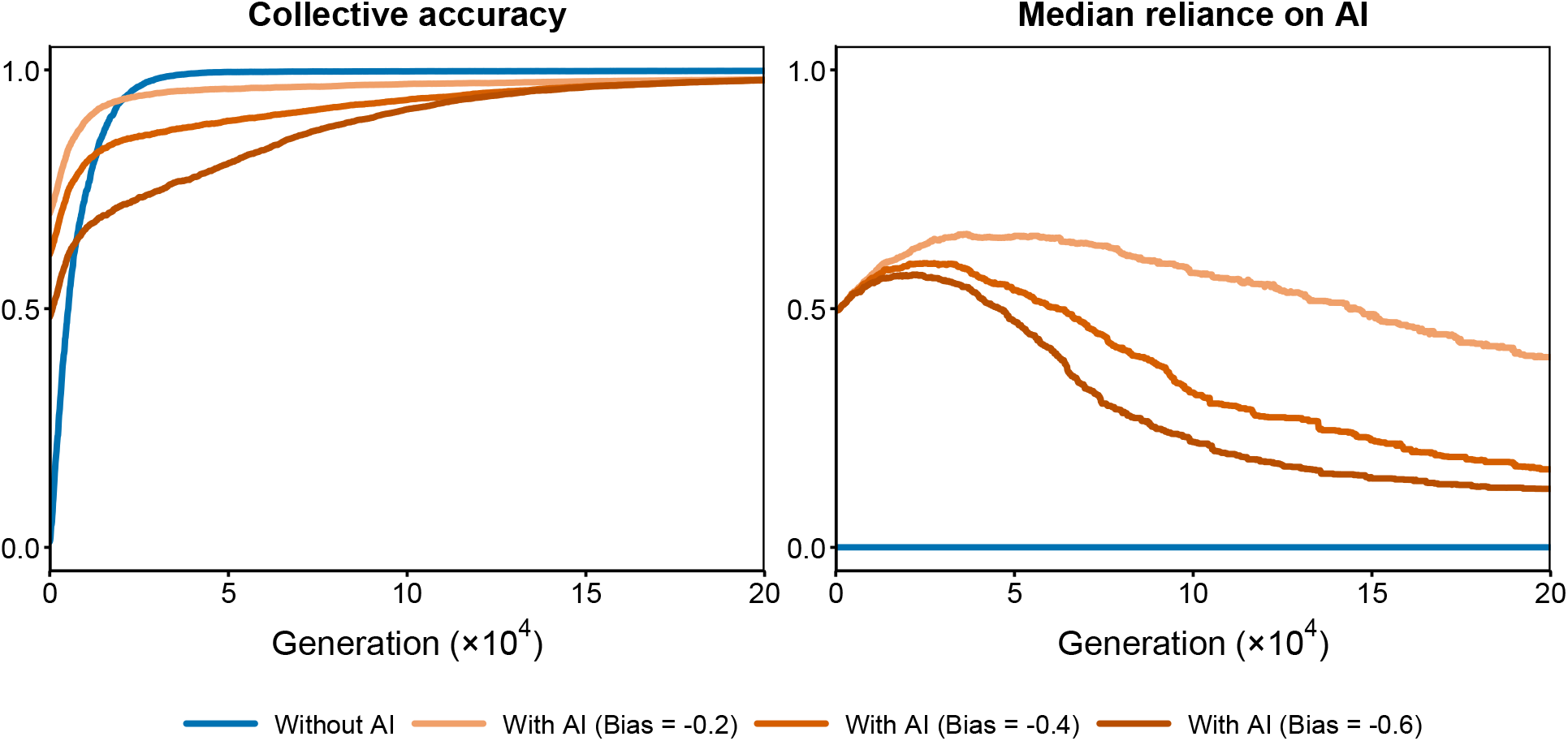
Niche-expert structure may reduce reliance on AI, delaying the emergence of CI. Trajectories of collective accuracy and median reliance on AI when the niche-expert structure causes reliance on AI to decline over time. Red lines represent trajectories with AI across varying levels of AI bias (-0.2, -0.4, and -0.6). Green lines represent trajectories without AI; reliance on AI is set to zero in the absence of AI. This reduction in reliance on AI occurs when the *α*_0_ and AI bias have opposite signs, and is associated with the eventual emergence of CI, although CI emerges substantially later than in the condition without AI. Prediction target parameters: *M* = 50, *s* = 50, *µ* = 0, *α*_*i*_ ∼ *U* (−5, 5), and *σ*_*i*_ ∼ *U* (0, 3). Population parameters: *N* = 10,000, *c*_*A*_ ∼ *N* (0, 5^2^), and *β*_*A*_ ∼ *U* (0, 1). AI parameters: *τ* = 0.3, Chatbot AI is used.

**Figure S6:**
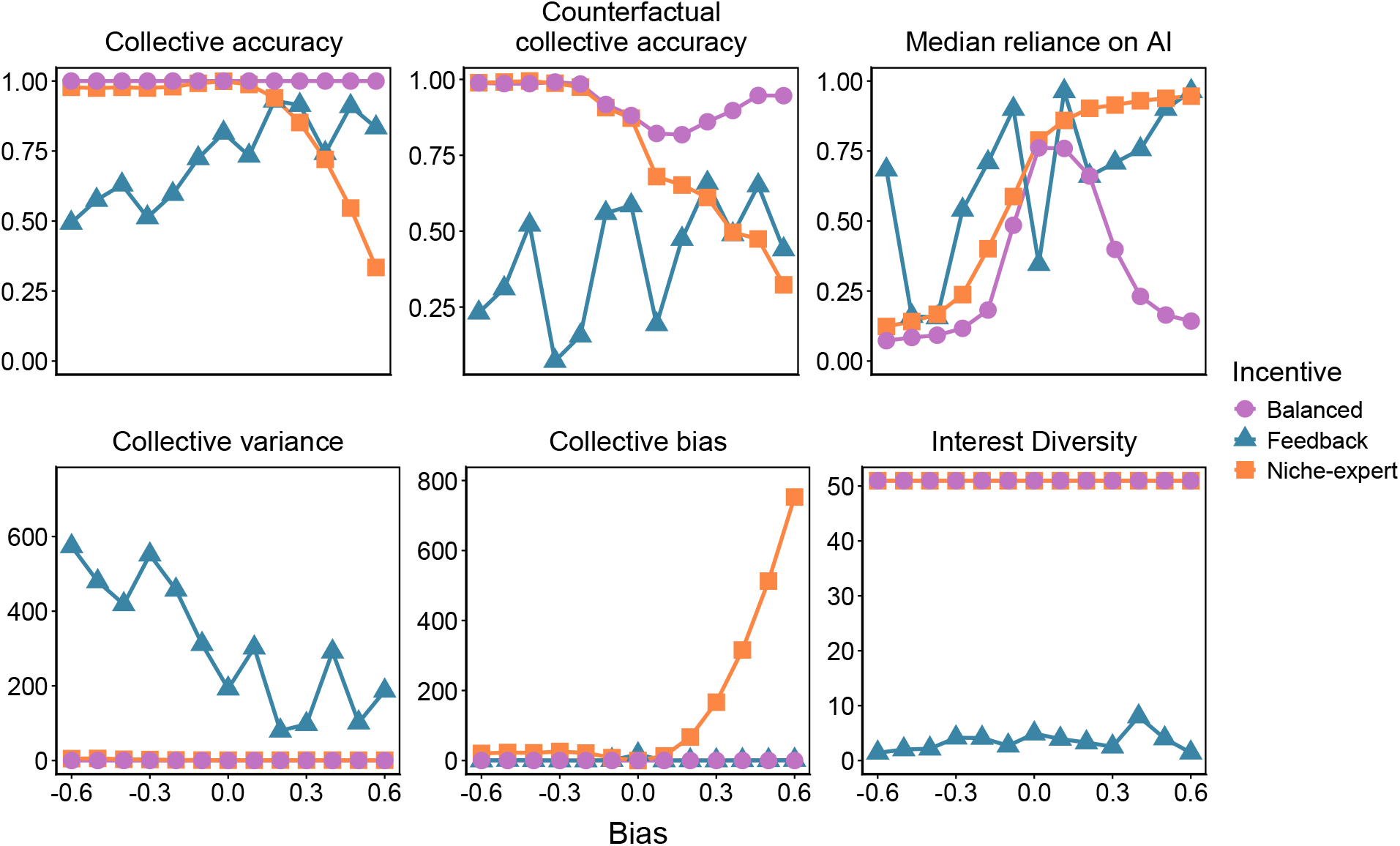
Comparison of evaluation metrics between incentive structures across various levels of AI bias. Evaluation metrics under feedback, niche-expert and balanced structures across varying levels of AI bias. Larger absolute AI bias indicates a less accurate AI model. All the evaluation metrics are averaged over generations 190,001–200,000. The balanced structure maintains near-perfect collective accuracy across bias levels and higher counterfactual human CI than the other incentive structures. These patterns are driven by well-calibrated reliance on AI, which increases as AI becomes more accurate, and by well-preserved interest diversity, which together lead to low collective variance and collective bias. Prediction target parameters: *M* = 50, *s* = 50, *µ* = 0, *α*_*i*_ ∼ *U* (−5, 5), and *σ*_*i*_ ∼ *U* (0, 3). Population parameters: *N* = 10,000, *c*_*A*_ ∼ *N* (0, 5^2^), and *β*_*A*_ ∼ *U* (0, 1). AI parameters: *τ* = 0.3, Chatbot AI is used.

**Figure S7:**
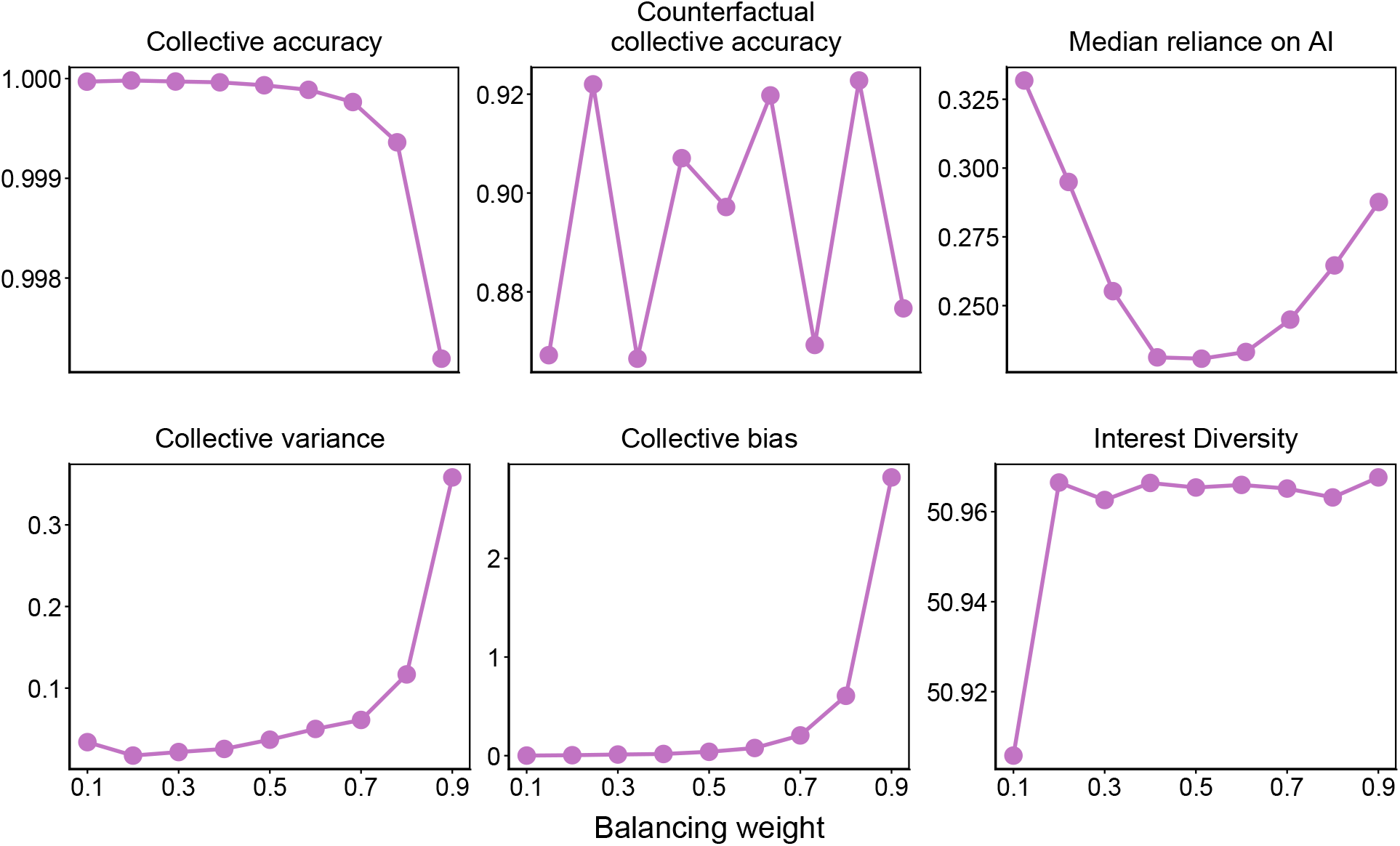
Balanced structure is robust across varying balancing weights. Evaluation metrics across varying levels of the balancing weight (0.1 – 0.9). The balancing weight represents the weight assigned to the niche-expert structure when constructing the balanced structure by combining the niche-expert and feedback structures. A balancing weight of 0.5 represents equal weighting of the niche-expert and feedback structures, whereas 0 corresponds to the feedback structure and 1 corresponds to the niche-expert structure. Although the evaluation metrics vary as the balancing weight changes from 0.1 to 0.9, the absolute magnitude of these deviations remains limited. Prediction target parameters: *M* = 50, *s* = 50, *µ* = 0, *α*_*i*_ ∼ *U* (−5, 5), and *σ*_*i*_ ∼ *U* (0, 3). Population parameters: *N* = 10,000, *c*_*A*_ ∼ *N* (0, 5^2^), and *β*_*A*_ ∼ *U* (0, 1). AI parameters: *b*_*i*_ = 0.4, *τ* = 0.3, Chatbot AI is used.

**Figure S8:**
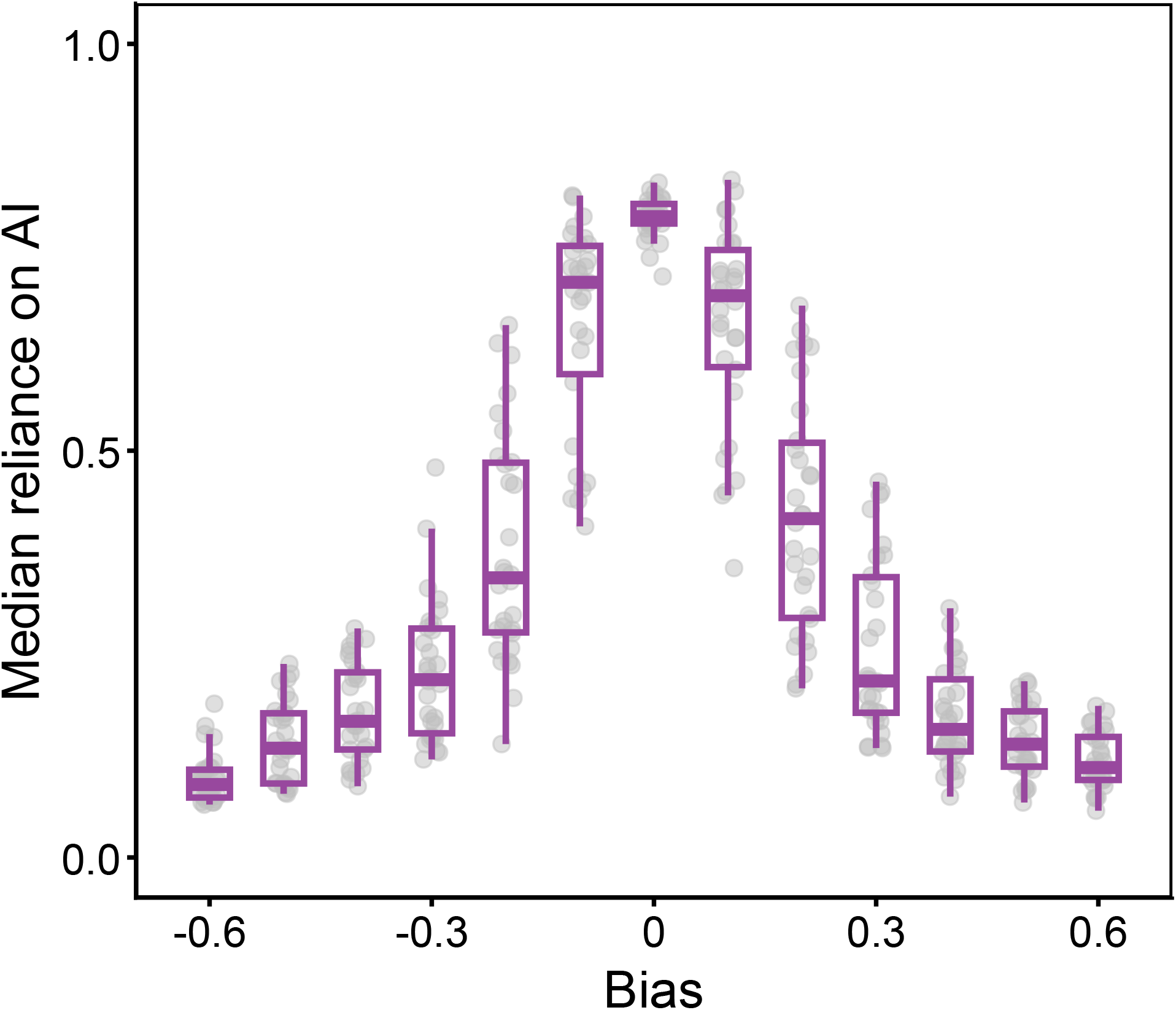
Balanced structure shows well-calibrated reliance on AI. Median reliance on AI across varying levels of AI bias. Thirty randomly initialized simulations were performed for each bias level. Each dot represents a single simulation, and the box plots show the median and interquartile range (IQR) for each bias level. The balanced structure shows well-calibrated reliance on AI, with reliance increasing as absolute magnitude of AI bias decreases. Prediction target parameters: *M* = 50, *s* = 50, *µ* = 0, *α*_*i*_ ∼ *U* (−5, 5), and *σ*_*i*_ ∼ *U* (0, 3). Population parameters: *N* = 10,000, *c*_*A*_ ∼ *N* (0, 5^2^), and *β*_*A*_ ∼ *U* (0, 1). AI parameters: *τ* = 0.3, Chatbot AI is used.

**Figure S9:**
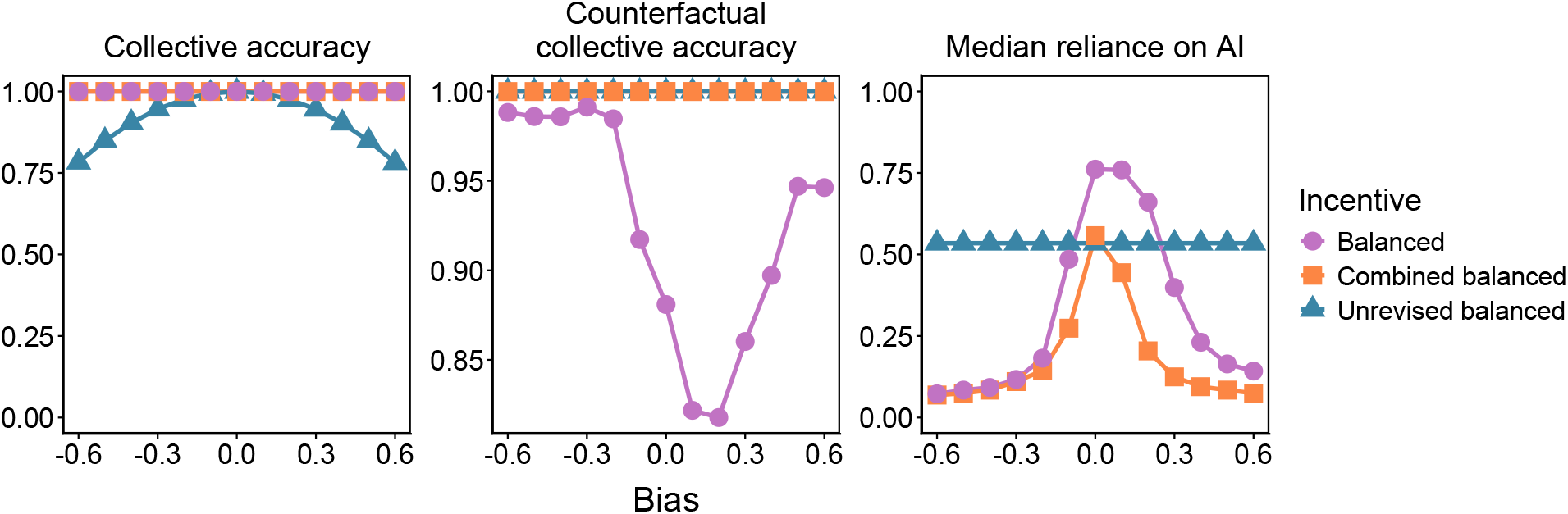
Incentive structures incorporating unrevised prediction accuracy can promote better counterfactual human CI. Evaluation metrics under the balanced, unrevised balanced, and combined balanced structures across varying levels of AI bias. The unrevised balanced structure is defined as the arithmetic mean of the feedback and niche-expert structures based on unrevised predictions, whereas the combined balanced structure is defined as the arithmetic mean of the balanced and unrevised balanced structures. All evaluation metrics are averaged over generations 190,001–200,000. The unrevised balanced structure achieves near-perfect counterfactual human CI; however, median reliance on AI remains near 0.5 because the unrevised balanced incentive does not account for AI predictions and therefore imposes no selection on *β*_*A*_. Consequently, collective accuracy directly inherits AI bias and decreases as the magnitude of AI bias increases. The combined balanced structure, which leverages both the balanced and unrevised balanced structures, achieves near-perfect counterfactual human CI without compromising collective accuracy. Prediction target parameters: *M* = 50, *s* = 50, *µ* = 0, *α*_*i*_ ∼ *U* (−5, 5), and *σ*_*i*_ ∼ *U* (0, 3). Population parameters: *N* = 10,000, *c*_*A*_ ∼ *N* (0, 5^2^), and *β*_*A*_ ∼ *U* (0, 1). AI parameters: *τ* = 0.3; Chatbot AI is used.

**Figure S10:**
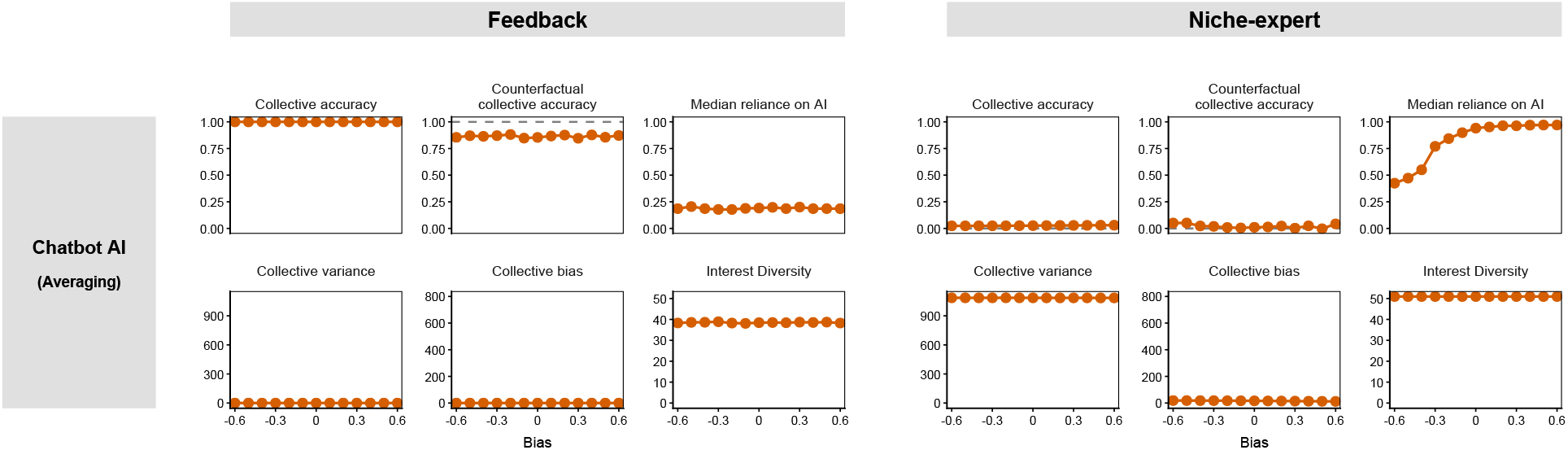
Chatbot AI with averaging aggregation has a limited influence on CI. Evaluation metrics across varying levels of AI bias under Chatbot AI with averaging aggregation. Two incentive structures, feedback and niche-expert, are considered. A larger absolute AI bias indicates a less accurate AI model. Dashed horizontal lines represent the values of the evaluation metrics when the group is not assisted by AI. Collective accuracy represents the predictive performance of the group, whereas counterfactual human CI represents the group’s predictive accuracy if AI assistance were unavailable at the given generation. Interest diversity is defined as the Hill–Shannon diversity of interests within the population, and median reliance on AI represents the median reliance on AI (*β*) across individuals. Collective bias represents the systematic deviation of the collective prediction from the true target, whereas collective variance represents the variability of collective prediction errors across prediction events. All evaluation metrics are averaged over generations 190,001–200,000. Prediction target parameters: *M* = 50, *s* = 50, *µ* = 0, *α*_*i*_ ∼ *U* (−5, 5), and *σ*_*i*_ ∼ *U* (0, 3). Population parameters: *N* = 10,000, *c*_*A*_ ∼ *N* (0, 100^2^), and *β*_*A*_ ∼ *U* (0, 1). AI parameters: *τ* = 0.3.

## Supplementary Table

**Table S1:**
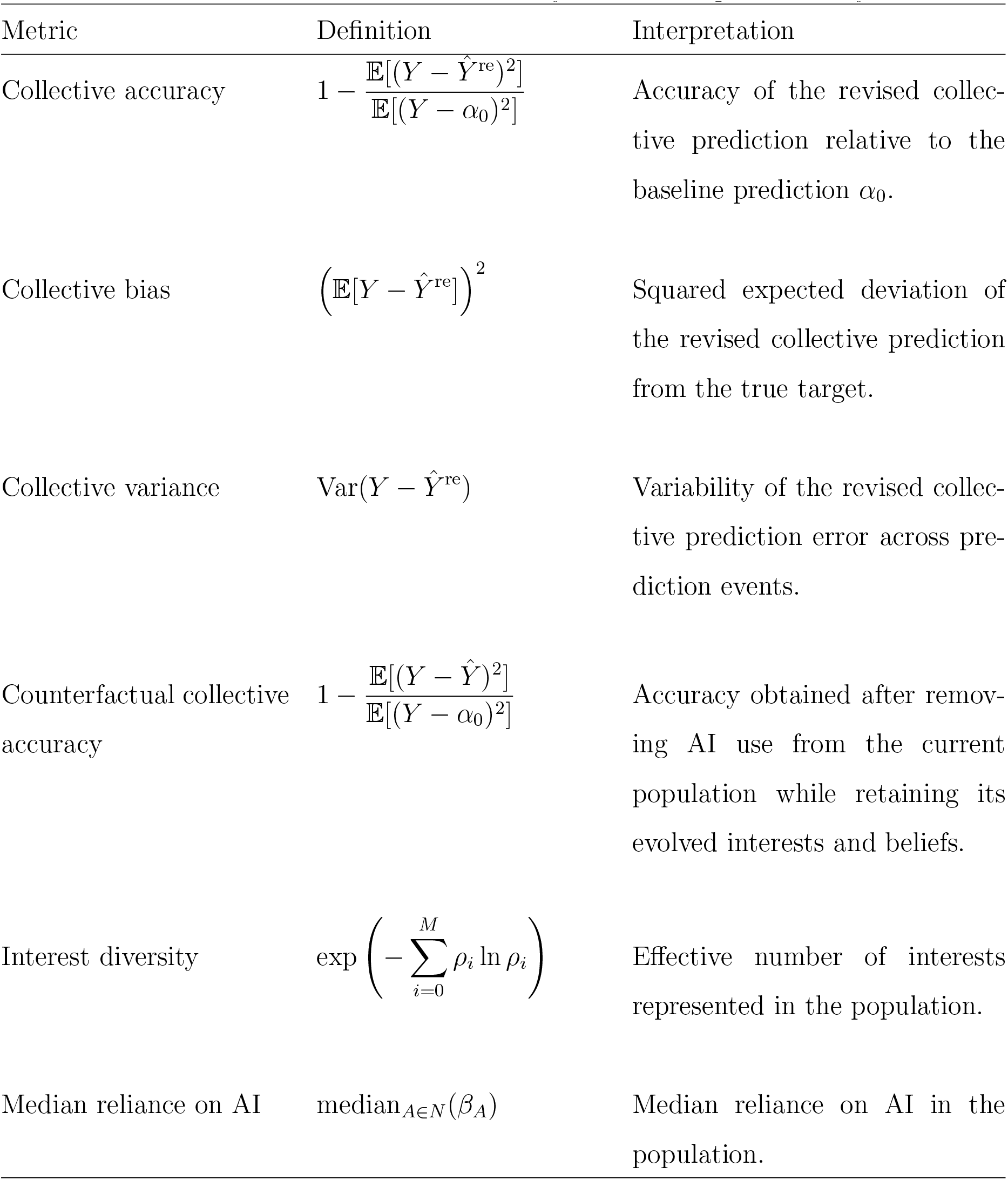
Evaluation metrics used to analyze collective prediction dynamics.

## References and Notes

1. F. Galton, Vox populi. Nature 75, 450–451 (1907).

2. I. Lorge, D. Fox, J. Davitz, M. Brenner, A survey of studies contrasting the quality of group performance and individual performance, 1920-1957. Psychological Bulletin 55 (6), 337 (1958).

3. I. Yaniv, M. Milyavsky, Using advice from multiple sources to revise and improve judgments. Organizational Behavior and Human Decision Processes 103 (1), 104–120 (2007).

4. A. W. Woolley, C. F. Chabris, A. Pentland, N. Hashmi, T. W. Malone, Evidence for a collective intelligence factor in the performance of human groups. Science 330 (6004), 686–688 (2010).

5. C. Hommes, J. Sonnemans, J. Tuinstra, H. Van de Velden, Coordination of expectations in asset pricing experiments. The Review of Financial Studies 18 (3), 955–980 (2005).

6. T. Dreyer, et al., Comparing cooperative geometric puzzle solving in ants versus humans. Proceedings of the National Academy of Sciences 122 (1), e2414274121 (2025).

7. A. M. Simons, Many wrongs: the advantage of group navigation. Trends in Ecology & Evolution 19 (9), 453–455 (2004).

8. A. Berdahl, C. J. Torney, C. C. Ioannou, J. J. Faria, I. D. Couzin, Emergent sensing of complex environments by mobile animal groups. Science 339 (6119), 574–576 (2013).

9. C. Riedl, Y. J. Kim, P. Gupta, T. W. Malone, A. W. Woolley, Quantifying collective intelligence in human groups. Proceedings of the National Academy of Sciences 118 (21), e2005737118 (2021).

10. L. Breiman, Bagging predictors. Machine Learning 24 (2), 123–140 (1996).

11. L. Breiman, Random forests. Machine Learning 45 (1), 5–32 (2001).

12. R. P. Mann, D. Helbing, Optimal incentives for collective intelligence. Proceedings of the National Academy of Sciences 114 (20), 5077–5082 (2017).

13. G. Wang, Q. Su, L. Wang, J. B. Plotkin, Individual incentives that promote collective intelligence. Proceedings of the National Academy of Sciences 122 (51), e2516535122 (2025).

14. J. Werfel, Fluid thinking about collective intelligence. Nature Machine Intelligence 8 (4), 506–516 (2026).

15. H. Cui, T. Yasseri, AI-enhanced collective intelligence. Patterns 5 (11), 101074 (2024).

16. S. Noy, W. Zhang, Experimental evidence on the productivity effects of generative artificial intelligence. Science 381 (6654), 187–192 (2023).

17. E. Brynjolfsson, D. Li, L. Raymond, Generative AI at work. The Quarterly Journal of Economics 140 (2), 889–942 (2025).

18. C. Lu, et al., Towards end-to-end automation of AI research. Nature 651 (8107), 914–919 (2026).

19. A. R. Doshi, O. P. Hauser, Generative AI enhances individual creativity but reduces the collective diversity of novel content. Science Advances 10 (28), eadn5290 (2024).

20. A. Bick, A. Blandin, D. J. Deming, The rapid adoption of generative AI. Management Science (2026).

21. S. Sajadieh, et al., Artificial Intelligence Index Report 2026. arXiv (2026).

22. M. Vaccaro, A. Almaatouq, T. Malone, When combinations of humans and AI are useful: A systematic review and meta-analysis. Nature Human Behaviour 8 (12), 2293–2303 (2024).

23. J. W. Burton, et al., How large language models can reshape collective intelligence. Nature Human Behaviour 8 (9), 1643–1655 (2024).

24. M. H. Tessler, et al., AI can help humans find common ground in democratic deliberation. Science 386 (6719), eadq2852 (2024).

25. A. Fügener, J. Grahl, A. Gupta, W. Ketter, Will humans-in-the-loop become borgs? Merits and pitfalls of working with AI. MIS Quarterly 45 (3), 1527–1556 (2021).

26. B. C. Lee, J. Chung, An empirical investigation of the impact of ChatGPT on creativity. Nature Human Behaviour 8 (10), 1906–1914 (2024).

27. L. Meincke, G. Nave, C. Terwiesch, ChatGPT decreases idea diversity in brainstorming. Nature Human Behaviour 9 (6), 1107–1109 (2025).

28. L. Hong, S. E. Page, Groups of diverse problem solvers can outperform groups of highability problem solvers. Proceedings of the National Academy of Sciences 101 (46), 16385–16389 (2004).

29. J. Lorenz, H. Rauhut, F. Schweitzer, D. Helbing, How social influence can undermine the wisdom of crowd effect. Proceedings of the National Academy of Sciences 108 (22), 9020–9025 (2011).

30. L. M. Aplin, D. R. Farine, R. P. Mann, B. C. Sheldon, Individual-level personality influences social foraging and collective behaviour in wild birds. Proceedings of the Royal Society B: Biological Sciences 281 (1789), 20141016 (2014).

31. J. Becker, D. Brackbill, D. Centola, Network dynamics of social influence in the wisdom of crowds. Proceedings of the National Academy of Sciences 114 (26), E5070–E5076 (2017).

32. I. Bousquette, Why Some Companies Say AI ‘Tokenmaxxing’ Is Key to Survival, https://www.wsj.com/cio-journal/why-some-companies-say-ai-tokenmaxxing-is-key-to-survival-e699a128 (2026), the Wall Street Journal. Accessed: 2026-07-07.

33. T. Keary, Is the cult of ‘Tokenmaxxing’ just another fad or the new normal?, https://www.forbes.com/sites/timkeary/2026/04/13/is-the-cult-of-tokenmaxxingjust-another-fad-or-the-new-normal/(2026), forbes. Accessed: 2026-07-07.

34. H. H. Thorp, ChatGPT is fun, but not an author. Science 379 (6630), 313–313 (2023).

35. D. S. Chawla, Researchers who use hallucinated references to face arXiv ban. Nature 653 (8116), 988–989 (2026).

36. L. Hong, S. E. Page, M. Riolo, Incentives, information, and emergent collective accuracy. Managerial and Decision Economics 33 (5-6), 323–334 (2012).

37. E. M. Bender, T. Gebru, A. McMillan-Major, S. Shmitchell, On the dangers of stochastic parrots: Can language models be too big?, in Proceedings of the 2021 ACM conference on fairness, accountability, and transparency (2021), pp. 610–623.

38. Q. Hao, F. Xu, Y. Li, J. Evans, Artificial intelligence tools expand scientists’ impact but contract science’s focus. Nature 649 (8099), 1237–1243 (2026).

39. L. Vicente, H. Matute, Humans inherit artificial intelligence biases. Scientific Reports 13 (1), 15737 (2023).

40. Z. Buçinca, M. B. Malaya, K. Z. Gajos, To trust or to think: cognitive forcing functions can reduce overreliance on AI in AI-assisted decision-making. Proceedings of the ACM on Human-computer Interaction 5 (CSCW1), 1–21 (2021).

41. H. Bastani, et al., Generative AI without guardrails can harm learning: Evidence from high school mathematics. Proceedings of the National Academy of Sciences 122 (26), e2422633122 (2025).

42. S. Melumad, J. H. Yun, Experimental evidence of the effects of large language models versus web search on depth of learning. PNAS Nexus 4 (10), pgaf316 (2025).

43. Y. Ke, et al., AI-induced never-skilling in medical education. Nature Medicine 32, 1997–2006 (2026).

44. J. A. Reif, R. P. Larrick, J. B. Soll, Evidence of a social evaluation penalty for using AI. Proceedings of the National Academy of Sciences 122 (19), e2426766122 (2025).

45. L. Ouyang, et al., Training language models to follow instructions with human feedback. Advances in Neural Information Processing Systems 35, 27730–27744 (2022).

46. M. Glickman, T. Sharot, How human–AI feedback loops alter human perceptual, emotional and social judgements. Nature Human Behaviour 9 (2), 345–359 (2025).

47. P. D. Taylor, L. B. Jonker, Evolutionary stable strategies and game dynamics. Mathematical Biosciences 40 (1-2), 145–156 (1978).

48. A. Traulsen, J. C. Claussen, C. Hauert, Coevolutionary dynamics: from finite to infinite populations. Physical Review Letters 95 (23), 238701 (2005).

